# Awakening adult neural stem cells: NOX signalling as a positive regulator of quiescence to proliferation transition in the *Xenopus* retina

**DOI:** 10.1101/2022.11.06.515333

**Authors:** A. Donval, CV Hernandez Puente, A. Lainé, D. Roman, R. Vessely, J. Leclercq, M. Perron, M. Locker

## Abstract

Reactive oxygen species (ROS) are both harmful molecules sustaining the pathogenesis of several diseases and essential modulators of cell behaviours. In particular, a growing wealth of data suggest that ROS-dependent signalling pathways might be critical in conferring embryonic or adult stem cells their specific properties. However, how stem cells control ROS production and scavenging, and how ROS in turn contribute to stemness remain poorly understood. Using the *Xenopus* retina as a model system, we first investigated the redox status of retinal stem cells (RSCs). We discovered that they exhibit higher ROS levels compared to progenitors and retinal neurons and express a set of specific antioxidant genes. We next addressed the question of ROS functional involvement in these cells. Using pharmacological or genetic tools, we demonstrate that inhibition of NADPH oxidase (NOX)-dependent ROS production increases the proportion of quiescent RSCs. This is surprisingly accompanied by an apparent acceleration of the mean division speed within the remaining proliferating pool. Our data further unveil that such impact on RSC cell cycling is achieved by modulation of the Wnt/Hedgehog signalling balance. Altogether, we highlight that RSCs exhibit distinctive redox characteristics and exploit NOX signalling to limit quiescence and fine-tune their proliferation rate.

## Introduction

The intracellular redox state recently emerged as an intrinsic regulatory mechanism for adult and embryonic stem cell self-renewal, proliferation and differentiation (Bigarella et al. 2014; Sinenko et al. 2021). Among molecular species that contribute to the cellular redox status are reactive oxygen species (ROS) such as the superoxide ion (O_2·_-), hydrogen peroxide (H_2_O_2_), or the highly reactive hydroxyl radical (·OH). ROS are mainly produced in cells through the mitochondrial electron transport chain or the activity of NADPH oxidase complexes (NOX1-5 and DUOX I-II). Under normal physiological conditions, the generation of ROS is tightly regulated by a ROS scavenging system, ensuring redox homeostasis. This system includes enzymes such as catalases, superoxide dismutases, glutathione reductases/peroxidases, thioredoxins or peroxiredoxins (Marengo et al. 2016). Oxidative stress occurs when ROS generation outpaces the antioxidant machinery. The resulting accumulation of high levels of oxygen free radicals and radical-derived species leads to premature aging, cell death or cancer through oxidative damage of cellular components including DNA. Despite being harmful, ROS also constitute essential modulators of cell behaviours. At basal levels, these molecules are indeed involved in reversible reduction-oxidation processes known to modulate key cellular signalling pathways (Bigarella et al. 2014; Marengo et al. 2016).

Deciphering how stem cells maintain their redox homeostasis and how ROS in turn impact stem cell activity is highly relevant in different fields from regenerative medicine to cancer therapy. With regard to oxidative stress protection, it is known that cancer stem cells exhibit an intracellular redox balance distinct from that of differentiated cancer cells, which makes them resistant to ROS-mediated cell killing (Kim et al. 2019). The molecular basis of such resistance is not well understood, but it is likely shared by embryonic and adult stem cells, and even proposed as an innate stemness feature (Madhavan et al. 2006; Guo et al. 2010). Among the prevailing assumptions to explain the lower vulnerability of stem cells to oxidative mutagenesis are stronger antioxidant defence mechanisms, their location in hypoxic niches and/or a glycolytic metabolism occurring even in the presence of oxygen (known as the Warburg effect). These features presumably contribute to the low ROS content reported in embryonic pluripotent stem cells or adult stem cells, such as hematopoietic or mesenchymal stem cells (Bigarella et al. 2014; Sinenko et al. 2021). In these cell types and their respective progeny, a gradual increase of ROS levels occurs along the differentiation process. This scheme might however not apply to all cycling cell populations. Neural stem cells of the mammalian brain were indeed found to exhibit higher ROS levels compared to progenitors in *ex-vivo* assays (Le Belle et al. 2011; Adusumilli et al. 2021). Besides, some studies suggest that ROS levels may vary depending on stem cell functional states (quiescent, primed, or activated). Here again, data from the literature do not support a unique redox rule for all stem cell types. In the hematopoietic system, low endogenous ROS content is associated with a greater quiescence (Jang and Sharkis 2007). In contrast, a study combining single-cell transcriptomic and ROS content labelling recently highlighted that quiescent neural stem cells from the dentate gyrus exhibit higher ROS levels compared to activated ones (Adusumilli et al. 2021).

From a functional point of view, fluctuations of ROS above or below their physiological levels were shown to affect a variety of processes including stem cell reprogramming (Zhou 2016), self-renewal, proliferation, and fate (Bigarella et al. 2014; Prozorovski et al. 2015; Rampon et al. 2018). However, somehow conflicting data were reported. For instance, increased ROS levels trigger a dramatic decline of hematopoietic stem cell number, associated with enhanced cycling and premature differentiation (Suda et al. 2011; Prieto-Bermejo et al. 2018). In contrast, high ROS levels are likely required for proper self-renewal of neural stem cells in the mammalian subventricular zone (Le Belle et al. 2011). Besides, despite an ever-increasing number of studies regarding ROS involvement in stem cell biology, few addressed their function *in vivo* and most of them relate to the regeneration field (Love et al. 2013; Gauron et al. 2013 Ferreira et al. 2016; Zhang et al. 2016; Hameed et al. 2015; Tao et al. 2016). We here sought to investigate redox status and ROS function *in vivo*, in adult neural stem cells of the *Xenopus* retina.

In contrast to the mammalian situation, the amphibian retina harbours continuously active retinal stem cells (RSCs) that sustain tissue growth throughout the animal life and can be further recruited following damage to regenerate lost cells (Hidalgo et al. 2014; Miyake and Araki 2014). The RSC niche is located in a region called the ciliary marginal zone (CMZ), whose spatial organization is well defined. Slow cycling multipotent stem cells reside in the most peripheral part of the CMZ, followed more centrally by transit amplifying progenitors and then by their postmitotic progeny (Perron et al. 1998). A recent study by Albadri and collaborators reported that H_2_O_2_ production is dynamically regulated during retinogenesis and within the zebrafish CMZ, contributing to the fine regulation of the balance between proliferation and differentiation (Albadri et al. 2019). The specific role of ROS in RSCs was however not addressed. We here first describe that RSCs are endowed with a peculiar redox status characterized by higher ROS levels compared to retinal progenitors. This is associated with the specific expression of several antioxidant genes. Second, our functional analyses reveal a crucial role of NOX as an homeostatic regulator of the proliferation rate within the post-embryonic stem cell pool, that limits the ratio of quiescent cells, while likely fine-tuning the mean division speed within the proliferative cohort. We further demonstrate that such function is mediated by modulation of the balance between the canonical Wnt pathway and Hedgehog signalling.

## Results

### Retinal stem cells of the post-embryonic CMZ exhibit higher ROS content than progenitor and differentiated retinal cells

Using a ratiometric sensor transgenic line, Albadri and collaborators recently showed that the embryonic zebrafish retina exhibit higher H_2_O_2_ concentrations in the proliferating peripheral margin (that includes the forming CMZ), compared with the differentiating central part of the tissue (Albadri et al. 2019). To assess whether such observation holds true in the *Xenopus* post-embryonic retina, we adapted a protocol set up by Owusu-Ansah *et al.* (Owusu-Ansah et al. 2008) to image ROS levels *in vivo*, using the fluorescent sensor dihydroethidium (DHE). We first tested DHE specificity by treating embryos 2 hours with rotenone (ROT), an inhibitor of the mitochondrial electron transport chain complex I, known to increase O_2·_- production (Li et al. 2003) (Figure 1A). As expected, ROT-treated embryos exhibited a significant increase in fluorescence intensity, suggesting that DHE staining indeed reflects intracellular ROS levels in living *Xenopus* embryos (Figure 1B, C). Of note, ROT treatment also enhanced the fluorescence of the ROS sensor CM-H_2_DCFDA (Supplementary Figure 1 A-C). We next analysed DHE fluorescence on retinal sections at the end of embryogenesis (Supplementary Figure 1 D). In line with zebrafish results, we found a significantly higher fluorescent signal within the CMZ compared to the differentiated neural retina (Supplementary Figure 1E, F). Of note, intensity within the CMZ could be enhanced by a 2-hour ROT treatment, confirming the specificity of the staining (Supplementary Figure 1F). Further quantitative analysis in these embryos revealed no difference in labelling between stem cells and progenitors (Supplementary Figure 1F). At post-embryonic stages, however, a peripheral to central gradient of fluorescence was clearly observed with a maximal intensity at the tip of the CMZ, where stem cells reside (Figure 1 D-F). These data show that, similarly to the zebrafish situation (Albadri et al. 2019), ROS levels are dynamically regulated within the *Xenopus* retina, both temporally (embryonic *versus* tadpole stages) and spatially (peripheral *versus* neural retina). They further highlight that post-embryonic neural stem cells exhibit a particularly high ROS content compared to progenitors or differentiated neurons.

**Figure 1.**
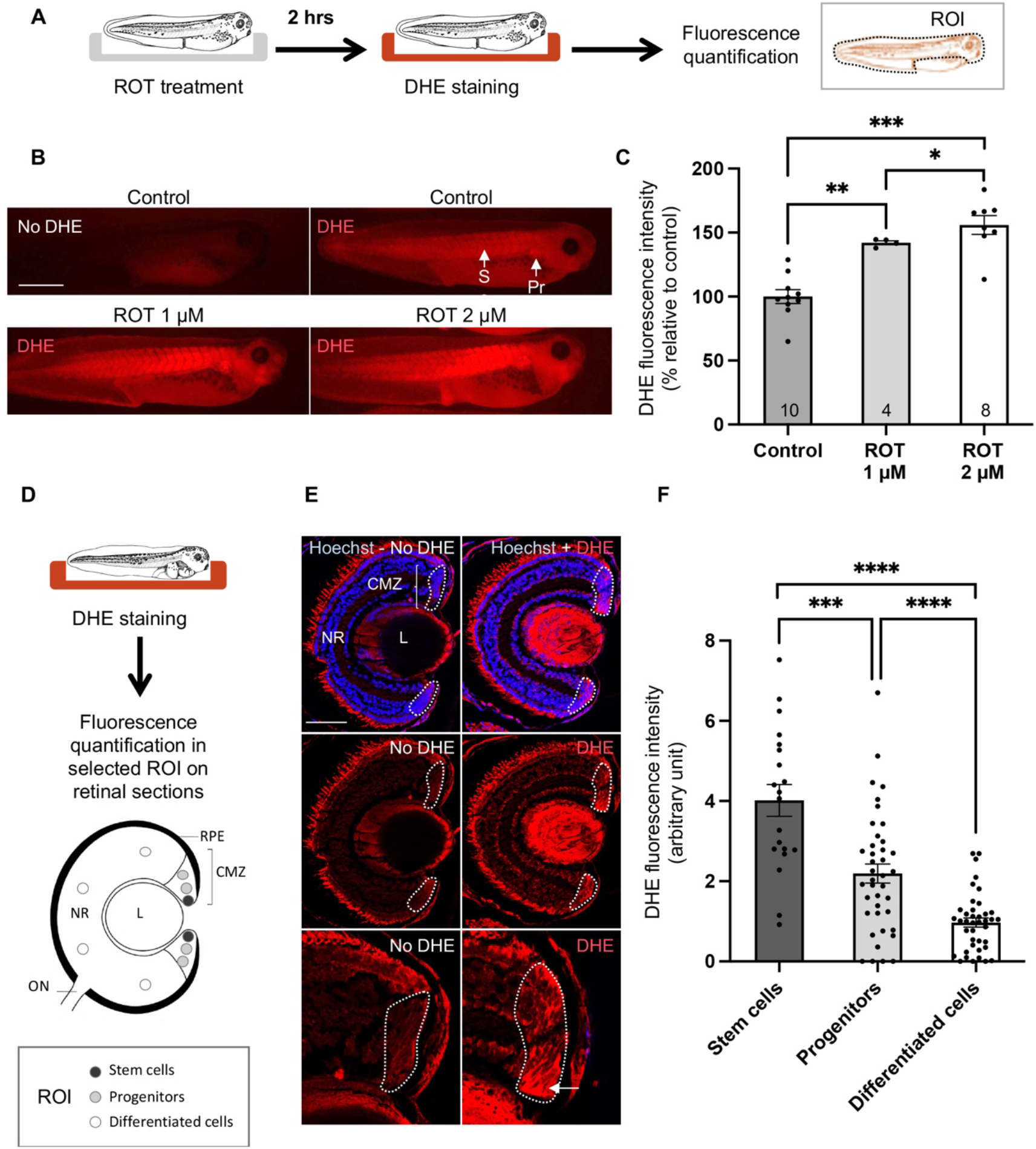
Redox status of post-embryonic RSCs. **(A)** Timeline diagram of the experimental procedure used in (B, C). Stage 39/40 control and rotenone (ROT)-treated tadpoles were labelled or not for ROS content with the fluorescent sensor dihydroethidium (DHE/no DHE). DHE intensity was quantified in the whole body (delineated in black). The vitellus was excluded because of its auto-fluorescence. **(B, C)** Corresponding images and quantifications. **(D)** Timeline diagram of the experimental procedure used in (E, F) to analyse DHE staining on retinal sections. Circles in the retina schematic indicate the 10 regions of interest (ROI) where DHE staining intensity was measured on each section. Dark and light grey circles are located in the stem cell and progenitor regions of the CMZ, respectively. White circles are located within the neural retina, where cells are differentiated. **(E)** Retinal sections from stage 42/43 tadpoles stained or not with DHE (DHE/no DHE). Nuclei were counterstained with Hoechst. Lower panels are higher magnifications of the dorsal CMZ (delineated in white). The white arrow points to RSCs. Note that in addition to the CMZ region, a strong DHE staining is detected within the lens, suggesting high ROS levels in this tissue as well. **(F)** Corresponding quantifications. In graphs, data are represented as mean ± SEM. In (C), the number of analyzed tadpoles is indicated in each bar. In (F), 10 sections were analyzed per condition. Statistics: Mann-Whitney test. CMZ: ciliary marginal zone; L: lens; NR: neural retina; ON: optic nerve; Pr: pronephros; RPE: retinal pigmented epithelium; So: somites. Scale bars: 1 mm in B and 50 μm in E.

### RSC express specific antioxidant genes

We next wondered whether such high ROS content might be associated with specific antioxidant defence mechanisms. In this purpose, we first cloned several *Xenopus* genes encoding antioxidant enzymes, including superoxide dismutases (*Sod1/2*), catalase 2 (*Cat2*), glutathione peroxidases 1/4/7 (*Gpx 1/4/7)* and peroxiredoxins 1-6 (*Prdx 1-6*). Superoxide dismutases are responsible for O_2·_- reduction into H_2_O_2_, while catalases, glutathione peroxidases and peroxiredoxins are involved in hydroperoxide detoxification. To our knowledge, the expression patterns of *Cat2, Sod1/2* and *Gpx1/4/7* genes were not described so far in developing *Xenopus laevis* embryos. Those of *Prdx* genes were already published but not analyzed on retinal sections (Shafer et al. 2011). Interestingly, none of the tested genes exhibited a uniform expression pattern but instead their mRNA proved to be enriched in very specific developing tissues and organs (Supplementary Figure 2). Of note, several were highly expressed in regions with high ROS levels (such as the somites or the pronephros), as inferred from DHE staining (Figure 1B). Regarding the eye (Figure 2A), *Gpx1* showed a lens-specific expression, while no obvious retinal signal could be detected for *Prdx5* and *Gpx7*. All the others were clearly expressed at the tip of the CMZ, as inferred from the ring-shaped labelling around the lens. Except for *Prdx2, Sod1* and *Sod2*, this was associated with a clear staining of the midbrain-hindbrain boundary, an expression pattern that we already described as typical of RSC markers (Parain et al. 2012). Further analysis on retinal sections did not allow to assess *Sod2* labelling (the signal was too faint) but confirmed that all the others were expressed in the stem cell-containing region (Figure 2B). However, their expansion within the CMZ significantly differed from one to another (Figure 2C). Three distinct profiles could be distinguished: an homogeneous expression in the CMZ (*Prdx2*), an enriched expression at the CMZ tip extending further at lower levels (*Prdx1*), or an expression strictly limited to the CMZ tip (encompassing stem cells and probably young progenitors; strong for *Cat2* and *Prdx3/6* but much lower for *Sod1, Gpx4* and *Prdx4*). As a whole, these results support the idea of distinct and tightly regulated antioxidant toolkits within the different CMZ domains and highlight *Cat2, Prdx3* and *Prdx6* as specific and highly expressed RSC markers.

**Figure 2.**
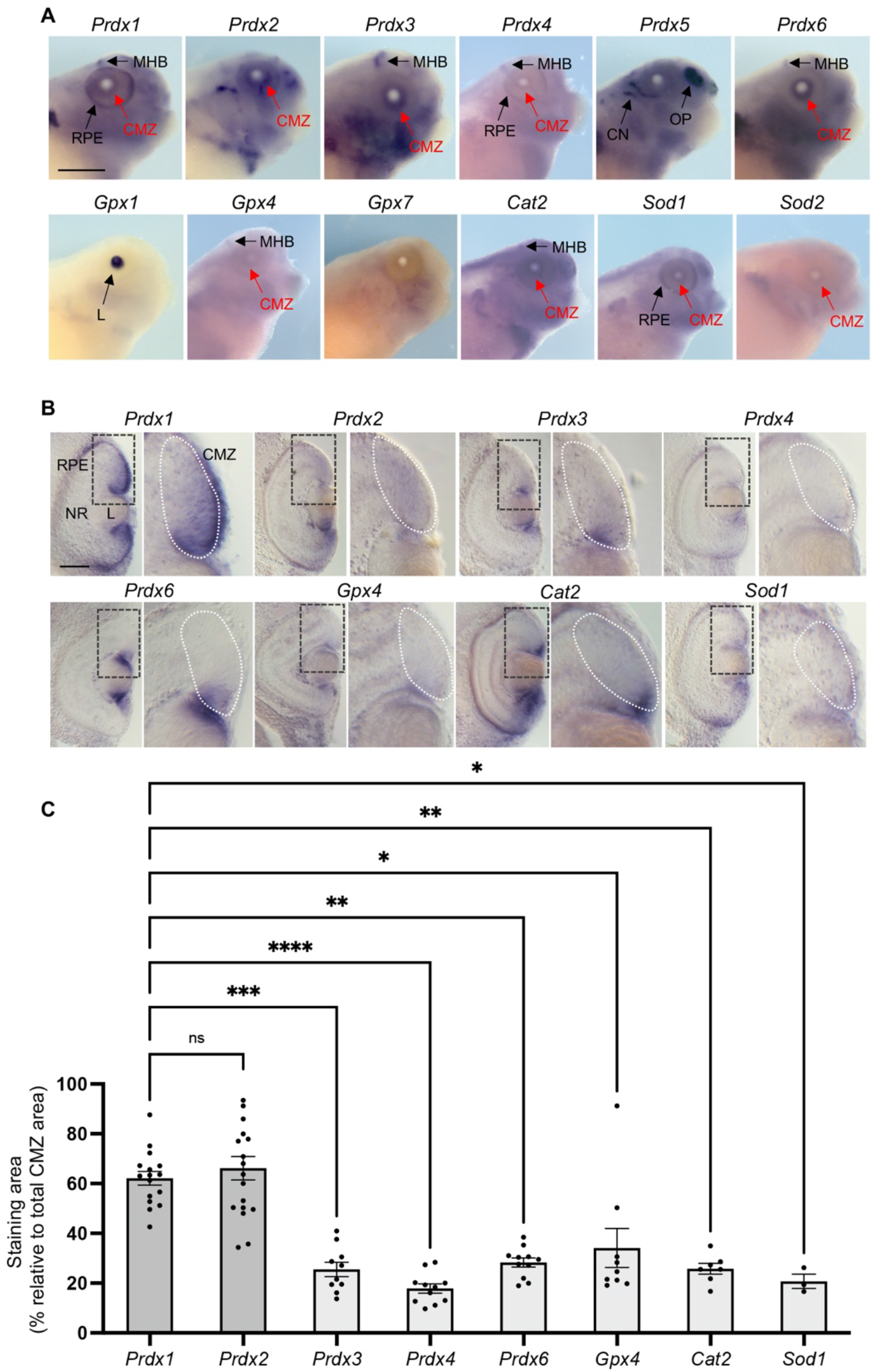
Identification of RSC specific antioxidant genes. **(A, B)** Whole mount *in situ* hybridization analysis of antioxidant gene expression on stage 38/39 embryos. Shown are representative images of stainings *in toto* (lateral views of the head; A) or on retinal sections (B). In B, right panels are higher magnifications of the dorsal CMZ (delineated in white). **(C)** Quantification of staining extension (peripheral to central) relative to total CMZ area (the dorsal CMZ only was considered). In the graph, each point represents one section. 7 to 18 vibratome sections (from 8 to 10 embryos) per probe were analyzed (only 3 for *Sod1*, whose signal was hardly detectable on most sections). Statistics: Kruskall-Wallis test (compared to *Prdx1*). Cat: catalase; CMZ: ciliary marginal zone; CN: cranial nerve; Gpx: glutathione peroxidase; L: lens; MHB: midbrain-hindbrain boundary; NR: neural retina; Prdx: peroxiredoxin; OP: olfactory placode; RPE: retinal pigmented epithelium; Sod: superoxide dismutase. Scale bars: 500 μm in A and 50 μm in B.

### NOX-dependent ROS signalling is required for proper RSC proliferative activity

The specific expression of antioxidant genes in RSCs suggests a tight control of redox homeostasis in these cells and thereby raises the question of ROS signalling functions with respect to stemness features. To address this issue, we first set up optimal conditions to reduce ROS production *in vivo*, using two pharmacological NOX inhibitors (diphenyleneiodonium, DPI and apocynin, APO) and the mitochondria-targeted ROS scavenger mitotempo (MITO). Working concentrations were chosen based on previous reports in *Xenopus* or zebrafish (Peterman et al. 2015; Love et al. 2013) and did not result in any obvious developmental defects (data not shown). To assess drug specificity and efficiency, treated tadpoles were stained with either DHE or CM-H_2_DCFDA (Supplementary Figure 3A). As expected, fluorescence of both sensors was significantly decreased following exposure to the pharmacological compounds (Supplementary Figure 3 B-E).

We then investigated the impact of lowering mitochondrial or NOX-dependent ROS production on CMZ cell proliferation. Tadpoles were treated with either MITO or APO for 16 hours and then subjected to an EdU incorporation assay (Figure 3A). We next determined the percentage of EdU-positive cells within the whole CMZ (mainly composed of progenitors) or among its most peripheral part, which include quiescent and self-renewing stem cells (Wan et al. 2016; Tang et al. 2017). MITO treatment did not affect EdU incorporation in either progenitors, or RSCs (Figure 3B, C). This suggests that CMZ cell proliferation is not highly dependent on mitochondrial ROS levels. APO treatment, however, generated a different result. Although the proportion of EdU-labelled cells was unchanged among progenitors, it was significantly reduced within RSCs compared to the control (Figure 3D, E). DPI treatment resulted in the same phenotype (Supplementary Figure 3F, G), suggesting distinct sensitivities of RSCs and progenitors to lowered NOX-derived ROS production.

**Figure 3.**
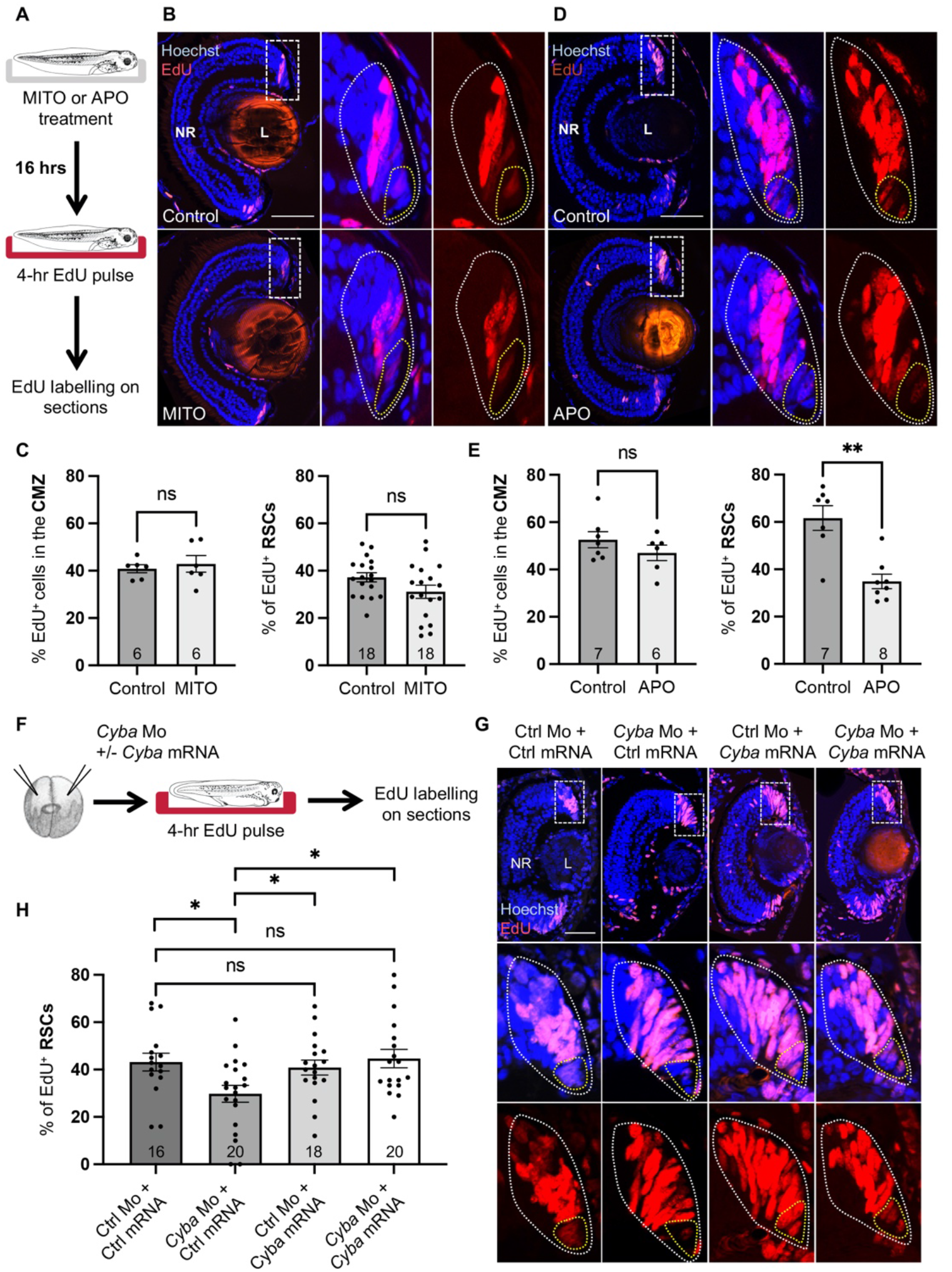
Impaired RSC proliferation upon NOX inhibition. **(A)** Timeline diagram of the experimental procedure used in (B-E). Tadpoles were treated for 16 hours with mitotempo (MITO; stage 43; B, C) or apocynin (APO; stage 41; D, E) and subjected to an EdU incorporation assay (4-hour exposure). (**B-E)** Representative images of retinal sections and corresponding quantifications. **(F)** Timeline diagram of the experimental procedure used in (G, H). Stage 37/38 embryos were subjected to an EdU incorporation assay (4-hour exposure), following co-injection at the two-cell stage of control (Ctrl) or *Cyba* Morpholinos (Mo), together with either control or *Cyba* mRNA (rescue construct insensitive to the morpholino). (**H, G)** Representative images of retinal sections and corresponding quantifications. In (B, D, G), nuclei were counterstained with Hoechst. Right panels in (B, D) or lower panels in (G) are higher magnifications of the dorsal CMZ (delineated in white). RSCs are delineated in yellow. EdUpositive cells were quantified among total CMZ cells or among RSCs. In graphs, data are represented as mean ± SEM. The number of analyzed retinas is indicated in each bar. Statistics: Mann-Whitney test. L: lens; NR: neural retina. Scale bars: 50 μm.

In order to further assess the absence of proliferative defects among progenitor cells, we examined the consequence of a longer APO-treatment on CMZ surface (as a reflect of total cell number) and neuronal production. Quantification of CMZ area following 4 days of APO exposure did not reveal any significant difference compared to the control (Supplementary Figure 4A, B). In addition, an EdU pulse-chase experiment revealed that the quantity of newborn cells produced by the CMZ over a period of a week was similar in APO-treated retinas and control ones (Supplementary Figure 4 C-E). This suggests that lowering NOX signalling does not impair the CMZ neurogenic activity, at least on the examined time-window.

We next aimed at confirming the effect of NOX inhibition by genetic means. In this purpose, we took advantage of a previously described translation-blocking Morpholino directed against *Cyba* mRNA (Love et al. 2013). The *Cyba* gene encodes a regulatory subunit in NOX complexes 1, 2 and 4, also called p22^Phox^. We first verified that this Morpholino was able to inhibit the translation of a Flag-tagged *Cyba* reporter construct (Supplementary Figure 5A, C). We next injected control or *Cyba* Morpholinos at the two-cell stage and analyzed EdU incorporation on retinal sections at the end of embryonic retinogenesis. Of note, *Cyba* knockdown frequently led to developmental defects affecting the ventral portion of the retina. We thus limited our quantification to the dorsal part of the CMZ. As observed with APO or DPI treatment, the proportion of EdU-labelled cells among progenitors was unchanged in Cyba morphant retinas compared to control ones (Supplementary Figure 5, E, F). The percentage of EdU-labelled RSCs was however significantly decreased (Figure 3 F-H). Importantly, the phenotype specificity was verified by a rescue experiment, in which the *Cyba* Morpholino was co-injected with a Morpholino-insensitive Myc-tagged *Cyba* mRNA (Figure 3 F-H and Supplementary Figure 5B, D). Altogether, these data demonstrate the requirement of NOX signalling for proper RSC proliferation. They also highlight the higher resilience of progenitors to reduced ROS production.

### Lowering NOX activity does not impair RSC survival and maintenance

We then wondered whether the defective proliferative phenotype of RSCs observed upon NOX inhibition might reflect their depletion. To address this question, we first assessed cell death following APO or DPI treatment. TUNEL assay (data not shown) or labelling against cleaved Caspase-3 did not reveal any significant increase of apoptotic cells in either the CMZ or the neural retina upon 16-hours of drug exposure (Supplementary Figure 6 A-C). After a 4-day APO treatment, a higher (although still limited) number of dying cells was found among treated progenitors and differentiated neurons compared to the control situation, showing some toxicity of the drug. However, here again, no apoptosis was detected within the RSC niche (Supplementary Figure 6 D-F). This shows that inhibiting NOX signalling does not impair RSC survival.

We next examined the expression of several RSC markers (*c-Myc, Prdx3, Hes1, Hes4* and *Yap)* by *in situ* hybridization after a 16-hour APO or DPI treatment (Figure 4A). We also included the examination of *Atoh7*, a marker of progenitors. None of them showed altered staining, reinforcing the view that progenitor cell number is not affected by reduced ROS levels, and suggesting that the RSC pool is conserved despite its lower proliferation rate (Figure 4B). To reinforce these data, we next extended APO exposure to 3 days, hypothesizing that a potential depletion of stem cells might take longer. Compared to a shorter treatment, such regimen did not aggravate the decreased EdU incorporation observed in RSCs (Figure 4C, D and compare with Figure 3E). In addition, qPCR analysis revealed no decrease of RSC marker expression. *Yap* was still properly expressed, and levels of *Hes1*, and to a lesser extent *Hes4*, were even found increased (Figure 4E, F). Taken together, these data suggest that NOX inhibition does not affect RSC maintenance.

**Figure 4.**
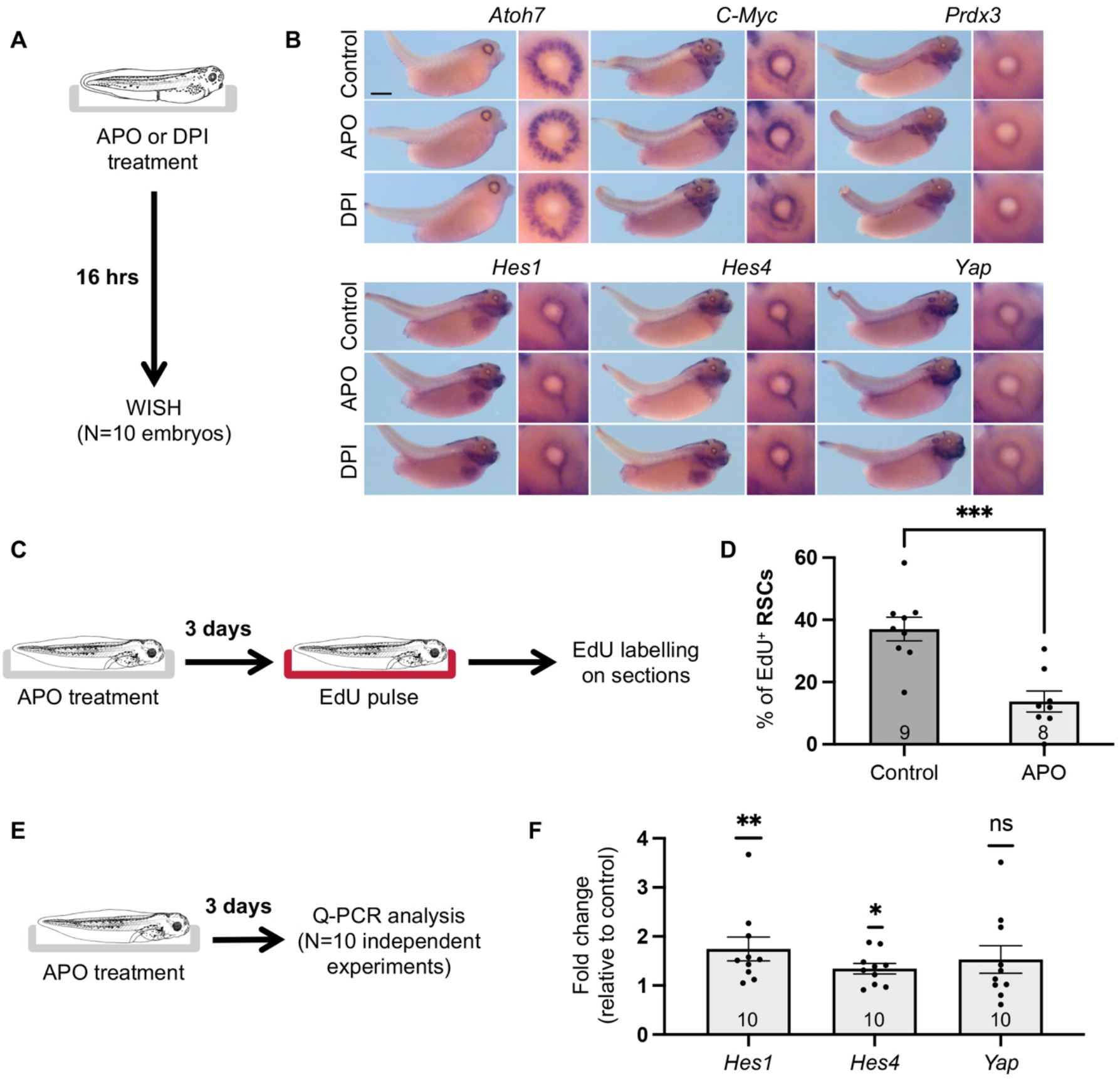
RSC marker expression upon NOX inhibition. **(A)** Timeline diagram of the experimental procedure used in (B). Stage 40 embryos were treated for 16 hours with apocynin (APO) or diphenyleneiodonium (DPI). They were then subjected to whole mount *in situ* hybridization (WISH) analysis of RSC (*c-Myc, Prdx3, Hes1, Hes4, Yap)* or progenitor (*Atoh7)* marker expression. **(B)** Representative images of stainings *in toto* (lateral views on the left and zoom on one eye on the right). **(C-F)** Analysis of RSC proliferation (C, D) and RSC marker expression (E, F) following a 3-day treatment with apocynin (APO) at stage 42/43. (C) and (E), are timeline diagrams of the experimental procedures used in (D) and (F), respectively. (D) EdU incorporation assay (4-hour exposure). (F) q-PCR analysis of *Hes1, Hes4* and *Yap* expression on eye extracts. Shown are fold changes relative to control (set to 1) for 10 independent experiments. In graphs, data are represented as mean ± SEM. The number of analyzed retinas (D) or cDNA samples (F) is indicated in each bar. Statistics: Mann-Whitney test (D) and Wilcoxon matched-pairs signed rank test (F). Scale bar: 1 mm.

### NOX inhibition enhances the proportion of quiescent RSCs

Interestingly, the transcriptional repressor Hes1 is known to actively maintain neural stem cell quiescence in the mouse brain (Sueda et al. 2019; Marinopoulou et al. 2021). The combination of increased *Hes1* expression and decreased EdU incorporation observed upon APO treatment led us to hypothesize that NOX inhibition might trigger RSC quiescence. To address this question, tadpoles were pre-treated with APO for 24 hours and then continuously exposed to EdU for a period of 2, 3 or to 7 days (Figure 5A). Such a protocol allows labelling all cycling cells as soon as the EdU pulse duration exceeds total cell cycle duration, thereby giving an evaluation of the growth fraction (GF; proportion of proliferating cells in a given population). The percentage of EdU-labelled RSCs was found stable in both control and APO-treated retinas at the three time-points, showing that the plateau was already reached after a 2 day-pulse. However, the labelling index was systematically and significantly reduced following APO treatment (Figure 5B, C). Such a decreased growth fraction reinforces the idea that inhibiting NOX signalling pushes RSCs into quiescence. Altogether, these data strongly suggest that NOX signalling is required to limit the proportion of quiescent RSCs within the CMZ.

**Figure 5.**
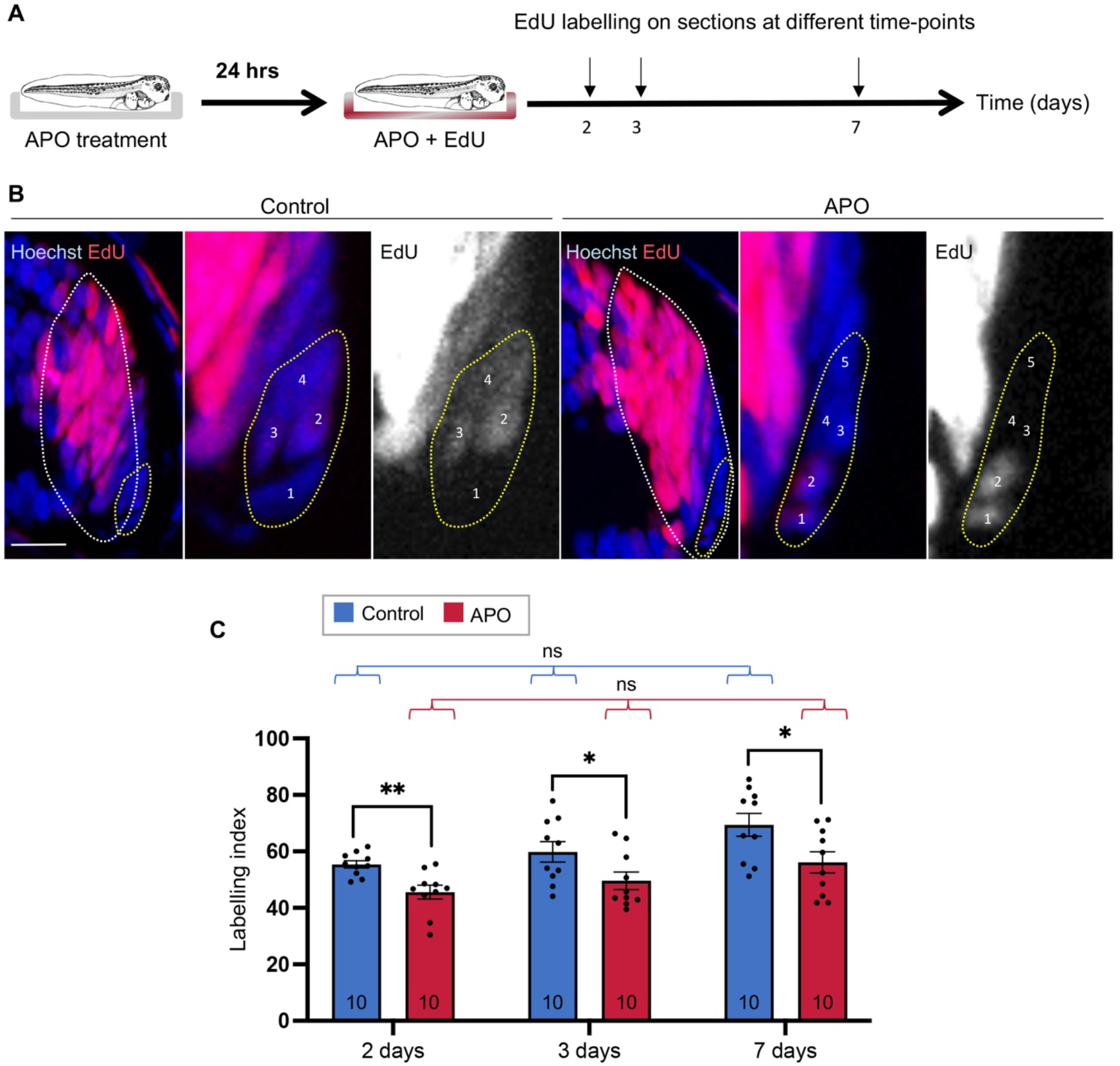
RSC entry into quiescence upon NOX inhibition. **(A)** Timeline diagram of the EdU cumulative labelling procedure used in (B, C). Stage 41/42 tadpoles were treated with apocynin (APO) for 24 hrs before addition of EdU to the rearing medium. They were then fixed and processed for EdU labelling after 2, 3 or 7 days of EdU exposure. Both the drug and the EdU were renewed daily. **(B)** Representative images of the dorsal CMZ (delineated in white) after a 3-day exposure to EdU, time at which all proliferative RSCs have already been labelled. Nuclei were counterstained with Hoechst. Right panels are higher magnifications of the RSC niche (delineated in yellow). In these images, 4-5 RSC nuclei are pointed (numbered from 1 to 4 or 5). **(C)** Quantification of the EdU cumulative labelling index within RSCs, along with increasing EdU exposure times. Data are represented as mean ± SEM. The number of analyzed retinas is indicated in each bar. Statistics: Mann-Whitney test (pairwise comparison of the labelling index distributions between control and APO-treated retinas at each time point) or Kruskal Wallis test (comparison of the labelling index distributions at all time-points among control or APO-treated retinas). Scale bar: 25 μm.

### NOX inhibition results in apparent acceleration of RSC division speed within the remaining proliferating

Since a subset of APO-treated RSCs still proliferate, we next examined their cell cycle kinetics by assaying cumulative labelling following 1 to 72 hours of EdU exposure (Figure 6A). This well-established technique allows to determine total cell cycle (T_C_) and S-phase (T_S_) durations (Locker and Perron 2019). Here again, the labelling index of APO-treated RSCs was consistently lower than that of control ones, in both the linear part of the curve and after the plateau was reached (Figure 6B). Confirming the previous results (Figure 5C), the growth fraction, estimated using the Nowakowski excel sheet (Nowakowski et al. 1989), dropped from 60% to 33% (Figure 6C). In addition, we found that the duration of G2 + M + G1 (T_C_-T_S_) was also decreased following APO treatment, as inferred from the time-point at which the labelling index reached the plateau (Figure 6B). Calculation of T_S_ and T_C_ revealed that both parameters were reduced compared to the control situation (−63% for T_C_ and −52% for T_S_; Figure 6C). These data thus show that APO-treated RSCs that remain proliferative exhibit enhanced cell cycle speed compared to the control pool of proliferative cells.

**Figure 6.**
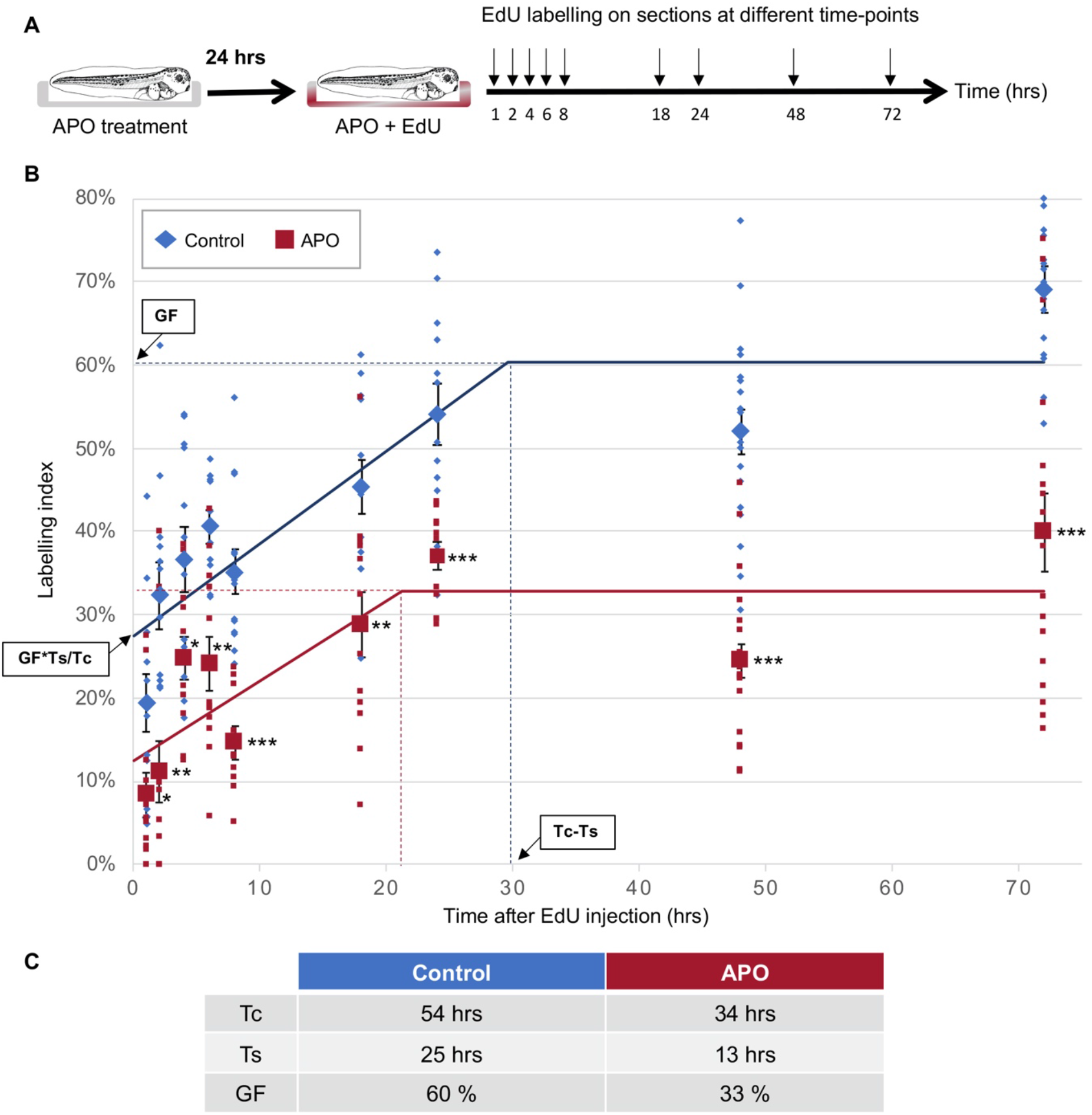
Enhanced cell cycle kinetics of RSCs upon NOX inhibition. **(A)** Timeline diagram of the EdU cumulative labelling procedure used in (B, C). Stage 41/42 tadpoles were treated with apocynin (APO) for 24 hrs before addition of EdU to the rearing medium. They were then fixed and processed for EdU labelling at different time points as indicated. Both the drug and the EdU were renewed daily. **(B)** Quantification of the EdU cumulative labelling index within RSCs, along with increasing EdU exposure times. The graph was made using the TcFit spreadsheet developed by R. Nowakowski, which allows drawing the best-fit line and calculates GF, T_C_, and T_S_ using a nonlinear regression method (Nowakowski et al. 1989). Small squares/lozenges represent 1 retina, while large ones correspond to mean ± SEM. Between 10 and 20 retinas were analyzed at each time-point and for each condition (corresponding to a total number of counted cells varying from 643 to 1173). Statistics: Mann-Whitney test. **(C)** Estimation of GF, T_C_ and T_S_ as calculated with the TcFit spreadsheet. GF: growth fraction; T_C_: total cell cycle length; T_S_: S-phase length.

### NOX-dependent ROS signalling regulates the Wnt/Hedgehog balance within the CMZ

In order to further understand the molecular mechanisms underlying NOX-dependent control of RSC proliferation, we assayed whether the canonical Wnt pathway might be impacted by APO or DPI treatment. We considered it a good candidate as we previously showed its requirement for RSC proliferative activity (Denayer et al. 2008; Borday et al. 2012) and because it was reported to be positively modulated by ROS in other cellular contexts (Rampon et al. 2018). In order to examine Wnt activity, we took advantage of a *Xenopus tropicalis* transgenic reporter line, in which *GFP* expression is driven by a synthetic promoter harbouring seven optimal binding sequences for LEF-1/TCF (Figure 7A; Tran and Vleminckx 2014; Borday et al. 2018). We first aimed at verifying in this species that NOX inhibition yields the same proliferative phenotype than the one observed in *Xenopus laevis* tadpoles. We found indeed that APO treatment reduced EdU incorporation in RSCs, while leaving progenitors unaffected (Supplementary Figure 7 A-C). We next exposed transgenic embryos to APO or DPI for 16 hours and analyzed *GFP* expression on retinal sections. Compared to controls, treated individuals exhibited a significantly reduced *GFP* staining within the CMZ. Quantification within the RSC niche revealed a similar decreased intensity (Figure 7 B-D and Supplementary Figure 7 D-F). To reinforce these data, we also assessed Wnt activity following injection of *Cyba* Morpholinos at the two-cell stage. As observed with pharmacological treatments, morphant retinas displayed lower levels of *GFP* expression compared to control ones (Figure 7 E-G). These data demonstrate that the Wnt pathway is positively modulated by NOX.

**Figure 7.**
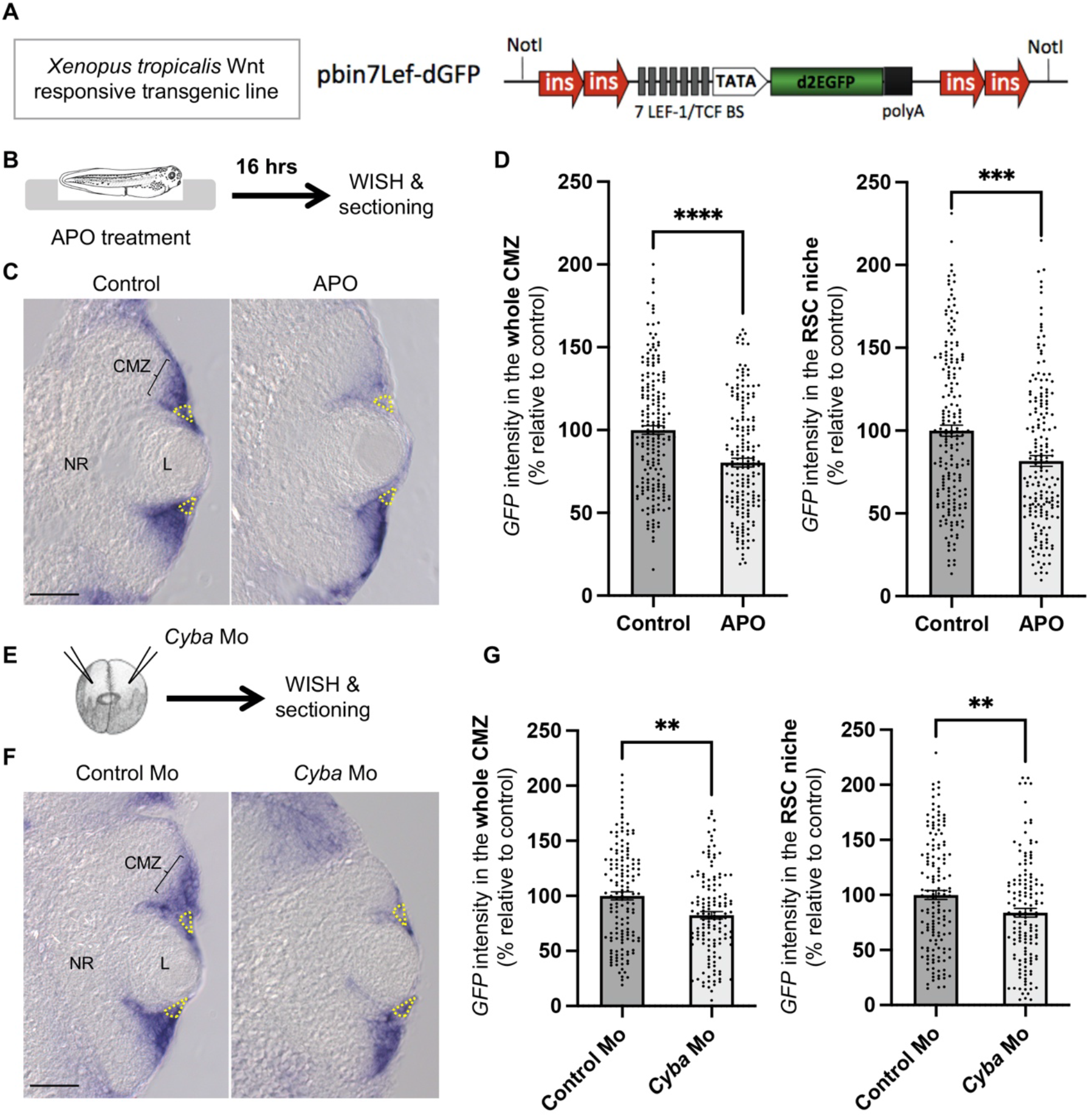
Decreased Wnt signalling activity upon NOX inhibition. **(A)** Schematic representation of the *pbin7Lef-dGFP* construct, which functions as a reporter of canonical Wnt signalling activity in transgenic (*Tg) Xenopus tropicalis.* **(B-G)** Whole mount *in situ* hybridization (WISH) analysis of *GFP* expression on retinal sections from stage 40 *Tg(pbin7Lef-dGFP)* embryos. Embryos were either treated for 16 hours with apocynin (APO; B-D) or injected at the two-cell stage with *Cyba* Morpholinos (Mo; EG). Shown in C, F are representative retinal sections. The RSC-containing region is delineated in yellow. In graphs, data are represented as mean ± SEM and each point corresponds to a measurement performed either in the dorsal or ventral part of the CMZ. 77 to 106 sections per condition were analyzed (from 18 to 21 embryos). Statistics: Mann-Whitney test. CMZ: ciliary marginal zone; L: lens; NR: neural retina. Scale bars: 50 μm.

We then investigated whether the decreased Wnt activity induced by NOX inhibition might be causal to the observed RSC proliferation defects. To investigate this question, we undertook a rescue experiment consisting in forcing Wnt activity in APO-treated embryos (Figure 8A). Canonical Wnt signalling activation was achieved by overexpressing an inducible and constitutively active form of TCF3 (TCF3-VP16GR; Borday et al. 2012), a transcriptional effector acting at the nuclear endpoint of the pathway. At the chosen injected dose, TCF-VP16GR by itself did not modify the proportion of EdU-labelled RSCs compared to the control situation. However, it efficiently rescued the decreased EdU incorporation observed upon APO treatment (Figure 8B, C). This highlights that NOX impact on RSC proliferation is mediated, at least in part, by Wnt signalling modulation.

**Figure 8.**
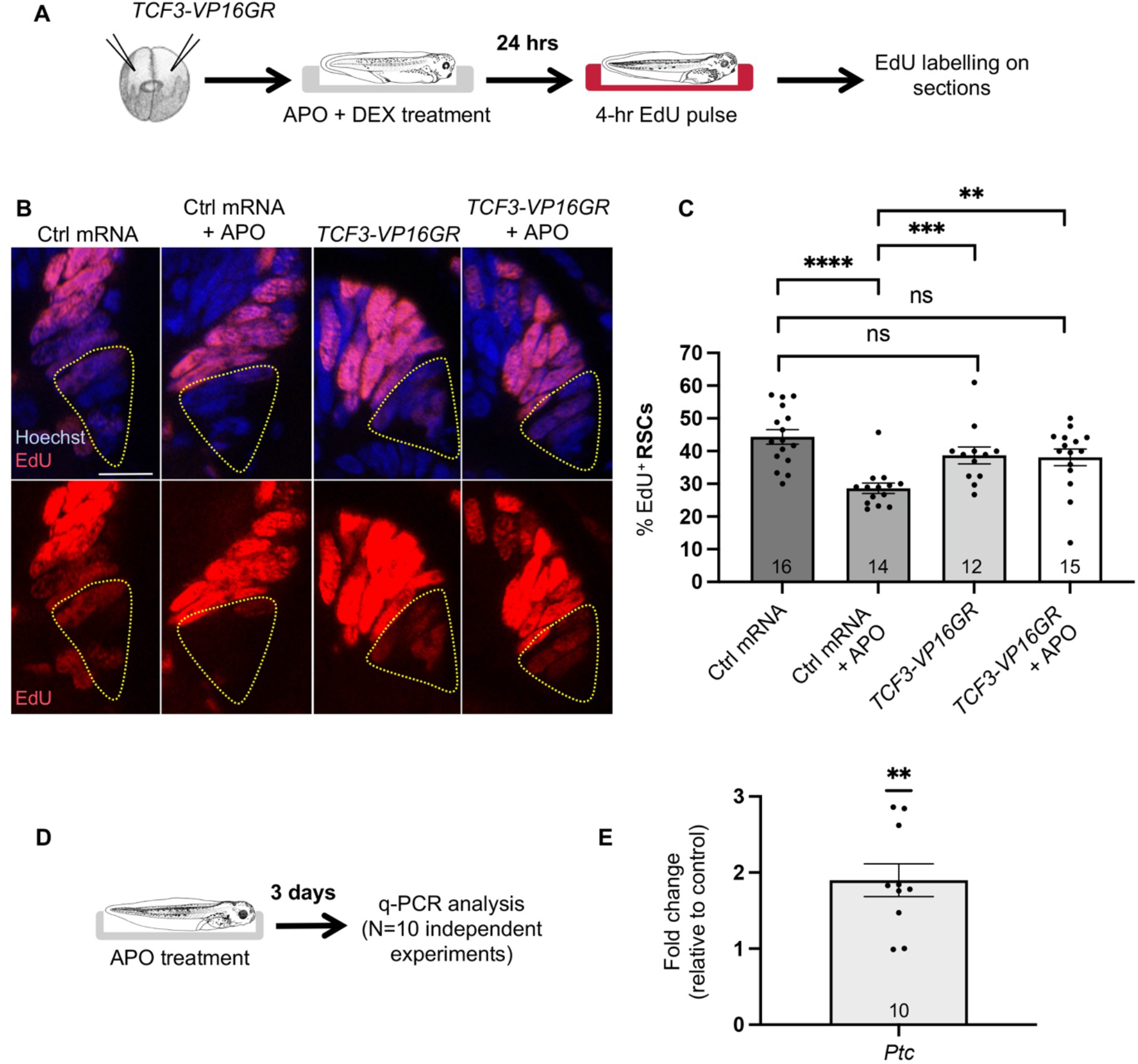
Functional interaction between NOX, Wnt and Hedgehog signalling. **(A)** Timeline diagram of the experimental procedure used in (B, C). Two-cell stage embryos were injected with control (Ctrl) or *TCF-VP16GR* mRNA. At stage 37/38, they were treated with dexamethasone (DEX; to induce TCF-VP16GR activity) and apocynin (APO) for 24 hours. They were then subjected to an EdU incorporation assay (4-hour exposure). (**B, C)** Representative images of retinal sections and corresponding quantifications. Nuclei were counterstained with Hoechst. RSCs are delineated in yellow. In the graph, data are represented as mean ± SEM. The number of analyzed retinas is indicated in each bar. Statistics: Mann-Whitney test. Scale bar: 25 μm. **(D)** Timeline diagram of the experimental procedure used in (E). Eye extracts were prepared from stage 42/43 tadpoles following a 3-day treatment with apocynin (APO). They were then subjected to q-PCR analysis of *Patched (Ptc)* expression. **(E)** Shown are fold changes relative to control (set to 1) for 10 independent experiments. Statistics: Wilcoxon matched-pairs signed rank test.

We have previously shown that CMZ cell proliferation is tightly regulated by the antagonistic interplay between Wnt and Hedgehog signalling (Borday et al. 2012). Since both pathways inhibit each other, we next addressed whether Hedgehog activity might be impacted as well upon NOX inhibition. In this purpose, we assayed expression of the Hedgehog target gene, *Patched.* We found it upregulated, as assessed by qPCR analysis (Figure 8D, E). NOX-dependent decrease of Wnt signalling is thus likely accompanied by enhanced Hedgehog activity.

Altogether, these data reveal that the Wnt/Hedgehog balance is regulated downstream NOX, and strongly suggest that modulation of these pathways account for ROS signalling effects on RSC proliferative activity.

## Discussion

Understanding how neural stem cells control their proliferative behaviour is a major issue for both regenerative medicine and cancer therapy. In this context, the amphibian retina model holds great advantage as it contains, like the fish, active neural stem cells, and is accessible to *in vivo* investigations. We here sought to decipher how the redox state might impact RSC activity. We first report that these cells exhibit high levels of ROS and express a specific set of antioxidant genes. Our functional investigations further reveal that RSCs exploit NOX-mediated ROS production to prevent their quiescence. Finally, we propose that such effect is mediated through regulation of the Wnt/Hedgehog signalling balance (Figure 9). Altogether, our study highlights NOX signalling as a novel player of the regulatory network that fine-tunes neural stem cell proliferative activity within the post-embryonic neurogenic niche of the retina.

**Figure 9.**
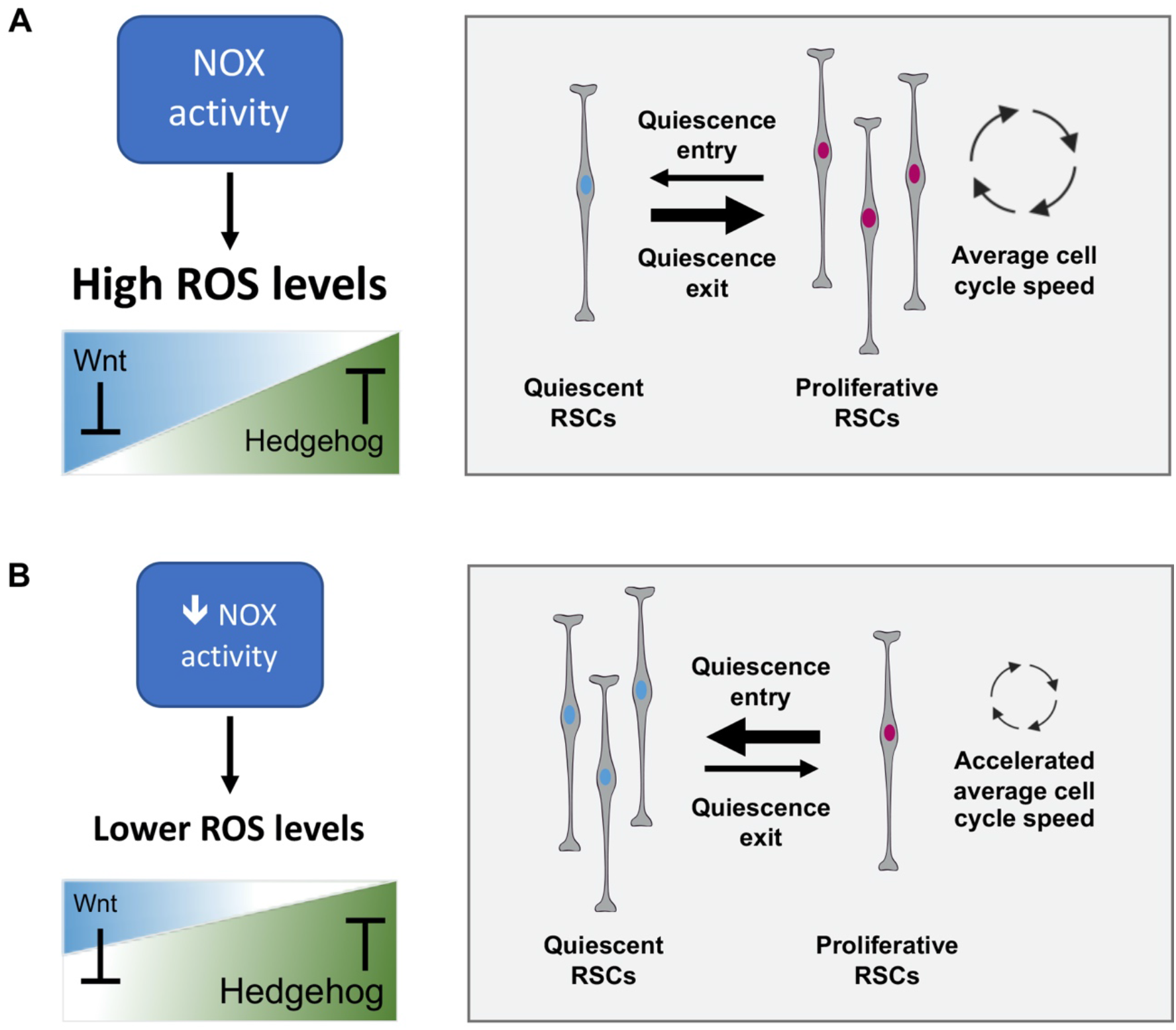
Model illustrating NOX signalling function in post-embryonic RCSs. **(A)** In the wild-type RSC niche, NOX-dependent control of the Wnt/Hedgehog balance results in low level of quiescence. Activated RSCs proliferate slowly. **(B)** Upon NOX inhibition, the activity of Wnt signalling drops, while that of the Hedgehog pathway increases. As a consequence, more RSCs are pushed towards quiescence. This is associated with an apparent increase of the mean cell cycle speed within the cell fraction that still proliferates.

Several studies have described that the route of neural stem cells towards terminal differentiation during development, is accompanied by an increase in ROS levels (due to a shift from glycolytic to oxidative metabolism), an enhancement of NOX activity, and an overall weakening of the antioxidant capacity (Olguín-Albuerne and Morán 2018). Contrasting with these data, Albadri and collaborators recently revealed in zebrafish that ROS levels are higher in the proliferating retinal margin compared to the differentiated neural retina (Albadri et al. 2019). We here confirm such peripheral to central gradient and further highlight that post-embryonic RSCs exhibit higher ROS content compared to neural progenitors. This is highly reminiscent of the adult mouse brain situation, where cells exhibiting the highest ROS level were found to correspond to self-renewing multipotent neural progenitors with phenotypic characteristics of neural stem cells (Le Belle et al. 2011; Adusumilli et al. 2021). Their finding and ours suggest that such an oxidative status constitutes a specific and conserved feature of adult neural stem cells. Associated with the observed high ROS content, we also found that several antioxidant genes are highly enriched or specifically expressed in RSCs. This is consistent with the general idea that stem cells exhibit stronger antioxidant defence. However, several antioxidant proteins, such as peroxiredoxins or thioredoxins, serve not only as guardians against oxidative stress but also, once oxidized by ROS, as mediators of redox signalling. Peroxiredoxin 6 for instance is a multitasking enzyme that modulates several pathways (Arevalo and Vázquez-Medina 2018) and was recently proposed to promote stem-like properties in small-cell lung cancer (Xu et al. 2019). The specific expression of several *Peroxiredoxin* genes in RSCs (in particular *Prdx3* and *Prdx6*) thus raises the question of their functions with regard to stemness properties.

If ROS involvement in neurogenesis is well-documented (Prozorovski et al. 2015; Bórquez et al. 2016; Wilson et al. 2018), how variations in their production mediate successive steps of proliferation and differentiation is still poorly understood (Bórquez et al. 2016). In addition, only a few studies have so far addressed the question of ROS functions in adult neural stem cells specifically. A great wealth of *in vivo* data indeed relates to embryonic precursors or, when conducted at the adult stage, do not truly distinguish bona fide stem cells from their more committed proliferative progeny (globally referring to neural stem/progenitor cells when examining for instance Sox2- or Nestin-positive cells). Both the subventricular zone and the dentate gyrus of the adult mammalian brain exhibit high expression levels of the NAPDH oxidase complex NOX2. Using knock-out mice, three different teams reported the requirement of NOX2-derived ROS to maintain normal proliferation and neuron production in these neurogenic regions (Dickinson et al. 2011; Le Belle et al. 2011; Nayernia et al. 2017). In these studies, however, whether and how the loss of NOX2 affects neural stem cell proliferative behaviour *in vivo* remains unclear. Only *ex vivo* data, using neurosphere assays, support the idea that ROS might be instrumental in neural stem cell selfrenewal (Paik et al. 2009; Le Belle et al. 2011). Taking advantage of the spatial organization of the CMZ, which allows for a topological distinction between stem cells and amplifying progenitors, we here identified a novel function of NOX-dependent ROS production in maintaining a proper ratio of activated neural stem cells within the CMZ. Further analyses should help decipher which intracellular or extracellular signals regulate NOX activity, and whether such a redox control of RSC proliferative activity might be coupled to changing tissue demand.

A functional link between low ROS levels and quiescence is known for long in hematopoietic stem cells (Kohli and Passegué 2014; Prieto-Bermejo et al. 2018). Current knowledge points to mitochondria, rather than to NOX enzymes, as the main source of ROS driving the transition of these cells to a proliferative state, as they escape their hypoxic niche and switch to a more energetic oxidative metabolism. In the developing brain, Khacho and collaborators recently revealed, that neural stem cells, although relying on glycolysis, possess functional elongated mitochondria, while committed progenitors exhibit fragmented ones. They further show that alteration of mitochondrial fusion/fission dynamics impairs the balance between self-renewal and commitment in a ROS-dependent manner (Khacho et al. 2016). Whether mitochondrial ROS production also contributes to fine-tune the proportion of proliferatively active neural stem cells is however still unknown. Our own data suggest that RSCs are not highly sensitive to decreased mitochondrial ROS production, with regard to their proliferative ability. We do not exclude, however, that longer treatment duration or higher concentration of the drug might provoke their dormancy and/or affect their maintenance or fate.

In addition to the increased rate of quiescence, we also surprisingly found that the mean division speed was enhanced upon NOX inhibition. A reasonable scenario to explain this phenotype might rely on a possible heterogeneity among RSCs in terms of cell cycle kinetics. With this respect, lowering NOX activity might push the more slowly proliferating ones to enter quiescence, leaving faster cells still active, and thus biasing the mean speed measure towards a higher value. Alternatively, NOX-derived ROS might directly act as a brake slowing down RSC divisions. Digging into the mechanisms that might underlie NOX impact on RSC proliferation, we found that NOX blockade results in reduced canonical Wnt signalling and enhanced Hedgehog activity. Furthermore, the defective RSC proliferation could be successfully rescued by forcing Wnt activation. These data thus raise two main questions: how does NOX regulate these pathways and how their regulation impacts RSC proliferative activity? We previously showed that both pathways are active in RSCs and reciprocally inhibit each other activity (Borday et al. 2012). NOX signalling could thus directly or indirectly modulate one, or the other, or both. Of note, NOX-derived ROS were already reported to modulate Wnt and Hedgehog signalling in diverse biological contexts (e.g. Love et al. 2013; Gauron et al. 2016; Thauvin et al. 2022), but little is known about the underlying molecular mechanisms. Only the Wnt pathway proved to be directly activatable by H_2_O_2_ through the oxidation of nucleoredoxin, a small redox-sensitive protein, which interacts with the adaptor protein Dishevelled (Funato et al. 2010). From a functional point of view, we previously showed that Hedgehog signalling has the ability to increase retinal precursor cell cycle kinetics (Locker et al. 2006). Although we did not examine its effect on RSCs specifically, its increased activity upon NOX inhibition might account for the apparent enhanced cell cycle speed observed. It might, however, also contribute to trigger RSC quiescence. In line with this idea, Daynac and collaborators examined the consequences of sustained Hedgehog signalling in adult neural stem cells of the subventricular zone through conditional deletion of its receptor Patched. They found a phenotype very similar to the one we observe in this study, with quiescent stem cells accumulating and the remaining active ones cycling faster (Daynac et al. 2016). In contrast to Hedgehog activation effects, increased canonical Wnt signalling (at least to a moderate level) proved sufficient to promote reactivation of hippocampal quiescent neural stem cells *in vitro* (Austin et al. 2021). The same was observed upon short-term *in vivo* clonal analyses of neural stem cells knocked-out in the *Sfrp3* gene (which encodes a secreted Wnt inhibitor) (Jang et al. 2013). Finally, our own data demonstrated a crucial role of Wnt signalling in the maintenance of CMZ cell proliferation (Denayer et al. 2008). Altogether, we thus propose that NOX activity is instrumental in controlling the rate of quiescent *versus* activated RSCs, and maybe their cell cycle speed, through its opposed effects on Wnt and Hedgehog activities.

## Materials and Methods

### Ethic statement

All animal cares and experimentations were conducted in accordance with institutional guidelines, under the institutional license C 91-471-102. The study protocol was approved by the institutional animal care committee, with the reference number APAFIS#5938-20160704l3l 04812 v2.

### Embryo collection, transgenic line

*Xenopus laevis* embryos and tadpoles were obtained by *in vitro* fertilization, staged according to Nieuwkoop and Faber (Nieuwkoop, P, Faber 1994) and raised at 18-21° C in 0.1X Modified Barth’s Saline (MBS). The *Wnt reporter transgenic Xenopus Tropicalis* line (carrying a *pbin7Lef-dGFP* transgene) was previously described and validated (Borday et al. 2018; Tran and Vleminckx 2014). Transgenic embryos were obtained by crossing a wild-type male with a homozygous transgenic female. They were grown in 0.05X Marc’s Modified Ringer (MMR) at 23°C.

### ROS detection and pharmacological treatments

The ROS sensor dihydroethidium (DHE, a reduced form of ethidium that is rapidly taken up by live cells and emits red fluorescence upon oxidation; ThermoFisher scientific, Waltham, MA, USA) was resuspended in 100 μL anhydrous DMSO to a 30 mM stock solution. The staining protocol was adapted from Owusu-Ansah and collaborators (Owusu-Ansah et al. 2008). Briefly, embryos/tadpoles were incubated in 0.1X MBS solution supplemented with a freshly made DHE solution (3 to 5 μM) for 10 minutes in complete darkness and then washed 3×5 minutes in 0.1X MBS. If previously treated with oxidant or antioxidant compounds, tadpoles were washed with 0.1X MBS solution supplemented with the drug they had been incubated in prior to DHE staining. The sensor 5-(and-6)-chloromethyl-2’,7’-dichlorodihydrofluorescein diacetate acetyl (CM-H_2_DCFDA, a chloromethyl derivative of H_2_DCFDA, whose oxidation yields a green fluorescent adduct that is trapped inside the cell; ThermoFisher scientific) was reconstituted in DMSO (2 mM stock solution). Tadpoles were incubated in darkness for two hours in MBS 0.1X containing 5 μM CM-H_2_DCFDA and then transferred for 30 min in MBS 0.1X to activate the probe. *In toto* imaging was performed immediately after, following tadpole anaesthesia in a 0.005% benzocaine solution (Sigma-Aldrich, Saint-Louis, MO, USA). Mitotempo stock solution (MITO, 3 mM; Sigma-Aldrich) was obtained by powder resuspension in dH_2_O and stored at −20°C. Apocynin (APO, 100 mM; Abcam, Cambridge, United Kingdom), diphenyleneiodonium (DPI, 600 μM; Sigma-Aldrich) and rotenone (ROT, 2 mM; Abcam) stock solutions were systematically made freshly following reconstitution in DMSO. These drugs were then added in the embryo rearing medium for various durations, as indicated, and at the following final concentrations: 20 μM MITO, 200 μM APO, 2-4 μM DPI, 1-2 μM ROT. The corresponding solvent was used as negative control.

### Plasmids and Morpholinos

cDNA encoding Catalase 2, Superoxide dismutases 1/2, Glutathione peroxidases 1/4/7, and Peroxiredoxins 1/2/3/4/5/6 were amplified by RT-PCR and inserted into a pCR®II-TOPO® vector through conventional TA-cloning procedures. PCR primer sequences are listed in Supplementary table 1. The *pCS2-TCF3-VP16GR* plasmid was described previously (Borday et al. 2012). pCS2+ plasmids encoding Myc- or Flag-tagged Cyba (*pCS2+-Myc-Cyba* and *pCS2+-Cyba-Flag)* were kindly provided by Enrique Amaya (Love et al. 2013). *Cyba* translation-blocking antisense Morpholino oligonucleotides (GeneTools, Philomath, OR, USA) were previously described and validated (Love et al. 2013). Of note, they perfectly matched with both the *Xenopus laevis* and *Xenopus tropicalis* target sequence. A standard Morpholino was used as a negative control (CCTCTTACCTCAGTTACAATTTATA).

### Microinjection

200 to 350 pg of mRNA (synthesized with mMessage mMachine kit; Life Technologies, Carlsbad, CA, USA) or 2 pmol Morpholinos were injected in both blastomeres at the two-cell stage. mRNAs encoding ß-Galactosidase and GFP were injected as controls and lineage tracers respectively. Activity of the TCF3-VP16GR chimeric protein was induced by incubating the embryos for 24 hours in 4 μg/ml dexamethasone (DEX, Sigma-Aldrich) from stage 37/38 to stage 41.

### Quantitative real-time PCR (qPCR)

Total RNA from 30 to 40 dissected eyes per condition was isolated using the Trizol reagent (Life Technologies). Reverse transcription was performed using the Superscript II enzyme (Invitrogen, Waltham, Massachusetts). qPCR reactions were performed in triplicate using SsoFast EvaGreen Supermix (BioRad, Hercules, CA, USA) on a CFX96 thermal cycler (BioRad). Results were normalized against the expression of reference genes *Odc* and *Rpl8.* Primer sequences are listed in Supplementary table 1.

### EdU labelling, immunostaining and western blot

Embryos/tadpoles were either injected intra-abdominally (at stage < 41) or bathed (at later stages) with/in a 1 mM 5-ethynyl-20-deoxyuridine solution (EdU, Invitrogen). They were then fixed after the desired time-period in 4% paraformaldehyde. Paraffin-embedded embryos/tadpoles were sectioned (11 μm) on a Microm HM 340E microtome (Thermo Scientific). EdU incorporation was detected on paraffin sections using the Click-iT EdU Imaging Kit according to manufacturer’s recommendations (Invitrogen). Cell nuclei were stained with Hoechst (Sigma-Aldrich). Immunostaining and western blots were performed with antibodies listed in Supplementary table 2, using standard procedures. Protein extracts were prepared using 8 embryos per condition (neurula stage), that were frozen-down on dry ice and then resuspended and homogenized in extraction buffer (20 mM Hepes, 15 mM MgCl_2_, 20 mM EDTA, 1mM DTT), supplemented with protease inhibitors. Extracts were centrifuged 10 min at 13000 rpm at 4°C and the interphase (devoid of pellets and lipids) was resuspended in migration buffer.

### Whole mount *in situ* hybridization

Digoxigenin-labelled antisense RNA probes were generated according to the manufacturer’s instructions (DIG RNA Labelling Mix; Roche, Bâle, Switzerland), from the corresponding plasmids previously linearized with the appropriate restriction enzymes. Whole mount *in situ* hybridization was carried out as previously described (Parain et al. 2012). Before imaging, embryos were post-fixed 20 min in 4% paraformaldehyde supplemented with 0.25% glutaraldehyde. Sections were either cut using a Leica VT1000S vibratome (50 μm thick) after 4% agarose embedding, or with a Microm HM 340E microtome (11 μm thick) following paraffin embedment.

### Imaging, quantification and statistical analysis

Fluorescence and brightfield images of whole mount embryos/tadpoles were captured with an AxioZoom fluorescence macroscope (Zeiss, Oberkochen, Germany). DHE or CM-H_2_DCFDA fluorescence intensity was quantified with Fiji software (Schindelin et al. 2012). Data were corrected by subtracting the mean fluorescence intensity of unstained negative controls and normalized to the corresponding ROI surface.

Retinal sections were imaged with an ApoTome-equiped Axio Imager.M2 microscope and processed using ZEN (Zeiss) and Photoshop CS5 (Adobe, Moutain view, CA, USA) softwares. Quantification of EdU-labelled cells was performed by manual counting in the CMZ. For each condition, 6 to 8 sections per retina were considered and a minimum of 6 retinas were analyzed. Stem cells were considered as the 4 to 6 most peripheral cells of the CMZ. EdU cumulative labelling experiments were analyzed as previously described (Locker and Perron 2019). Growth fraction (GF; proportion of proliferative cells), total cell cycle length (T_C_), and S-phase length (T_S_) were determined using the Excel spreadsheet provided by Dr R. Nowakowski (Nowakowski et al. 1989). All experiments were performed at least in duplicate. Shown in figures are results from one representative experiment. Unless indicated otherwise, statistical analyses were performed using the nonparametric Mann-Whitney test. Statistical significance is: ns: non-significant; *p < 0.05; **p < 0.01; ***p < 0.001; ****p < 0.0001.

## Acknowledgements

This research was supported by grants to MP and ML from the “Fondation ARC pour la Recherche sur le Cancer” and to MP from Retina France. CHP was granted by the Mexican government through a CONACYT fellowship. We thank Enrique Amaya for providing the Myc- and Flag-tagged Cyba plasmids.

## Author contributions

ML designed the work and wrote the manuscript. ML, AD, CHP, AL, DR, RV and JL carried out and analyzed experiments. MP analyzed experiments and reviewed the manuscript.

