## Supplementary Figures & Tables for "Awakening adult neural stem cells: NOX signalling as a positive regulator of quiescence to proliferation transition in the *Xenopus* retina"

| Amplicon | Cloning primers 5'-3' | Amplicon | q-PCR primers 5'-3' |
|---|---|---|---|
| ***Cat2*** | **F:** ATG GCG GAC AAG AGG GAT AAT | ***Odc*** | **F:** GCT TCT GGA GCG GGC AAA GGA |
| | **R:** CTA CAG GTT GGC TTT GTC CTT | | **R:** CCA AGC TCA GCC CCC ATG TCA |
| ***Sod1*** | **F:** ATG GTG AAG GCA GTG TGT GTG | ***Rpl8*** | **F:** CCA CGT GTC CGT GGT GTG GCTA |
| | **R:** TCA CGG ACT ATA ACC AAT CAC | | **R:** GCG CAG ACG ACC AGT ACG ACG A |
| ***Sod2*** | **F:** ATG CTG TGC AGG CTC AGT GTT | ***Hes1*** | **F:** AGC AAT ACC CCG GAT AAA CC |
| | **R:** TTA TTT TTT GGA AGC CTG ATA | | **R:** TCC AGG ATG AGG GTT TTG AG |
| ***Prdx1*** | **F:** ATG TCT GTC GGA AAT GCA AA | ***Hes4*** | **F:** CCC CCT CCA GCC AAC AAC CA |
| | **R:** TTA TTT CTG CTT GCT GAA GT | | **R:** GGG GGA GAT GGC CTC TGC TG |
| ***Prdx2*** | **F:** ATG GCG TGT CCT GTG CGT GC | ***Yap*** | **F:** GGC AAA GAC ACC CTC TGG GCA |
| | **R:** CGT TAG TAT TCT TTA GAG AAG A | | **R:** GGC GGG CTT GTG GGA GCA GTA |
| ***Prdx3*** | **F:** ATG GCG GCG TCC TGT GGA AG | ***Patched*** | **F:** CAG CTG CCC AGC CGA GGG TA |
| | **R:** TTA GTG CAC TTT CTC AAA GT | | **R:** GGG CGA AAT TGG CAT CGC AGT A |
| ***Prdx4*** | **F:** ATG GCT CTT CAG CTC CGG CG | | |
| | **R:** GTT TCA CTC CCA GGT TTC CA | | |
| ***Prdx5*** | **F:** ATG GCT CTT CGT ATC CCG GT | | |
| | **R:** CTT AAA GCT GAG ATA TTA TGT | | |
| ***Prdx6*** | **F:** ATG CCT GGA ATC CTG CTA GG | | |
| | **R:** TTA TTG TGG CTG TGC AGT GTA | | |
| ***Gpx1*** | **F:** ATG CGT TCA GCT ATG GTT TC | | |
| | **R:** CTA TGC ACA ATT TTC ATG AA | | |
| ***Gpx4*** | **F:** ATG TGT GCA CAA GCA GCA GA | | |
| | **R:** CTA CAG ATA ACT GGG TAG AT | | |
| ***Gpx7*** | **F:** ATG TAC CTT GCG GCT CTT GT | | |
| | **R:** TCA AAG TTC ATC TTT CTT TT | | |

**Supplementary table 1. List of primers used.**

| Antibody | Host | Supplier | Reference | Dilution IF | Dilution WB |
|---|---|---|---|---|---|
| **Anti-alpha Tubuline** | Mouse | Sigma | T5168 | | 1:50 000 |
| **Anti-Myc** | Mouse | proteintech | 60003-2-Ig | | 1:2000 |
| **Anti-FLAG** | Rabbit | SIGMA | F7425 | | 1:1000 |
| **HRP-conjugated anti-mouse** | Goat | Sigma-Aldrich | A4416 | | 1:5000 |
| **HRP-conjugated anti-rabbit** | Goat | Sigma-Aldrich | A0545 | | 1:5000 |
| **Anti-Cleaved Caspase-3** | Rabbit | Cell Signaling | 9661S | 1:250 | |
| **Alexa Fluor 594 anti-rabbit** | Goat | ThermoFisher Scientific | A11012 | 1:1000 | |

**Supplementary table 2. List of antibodies used.**

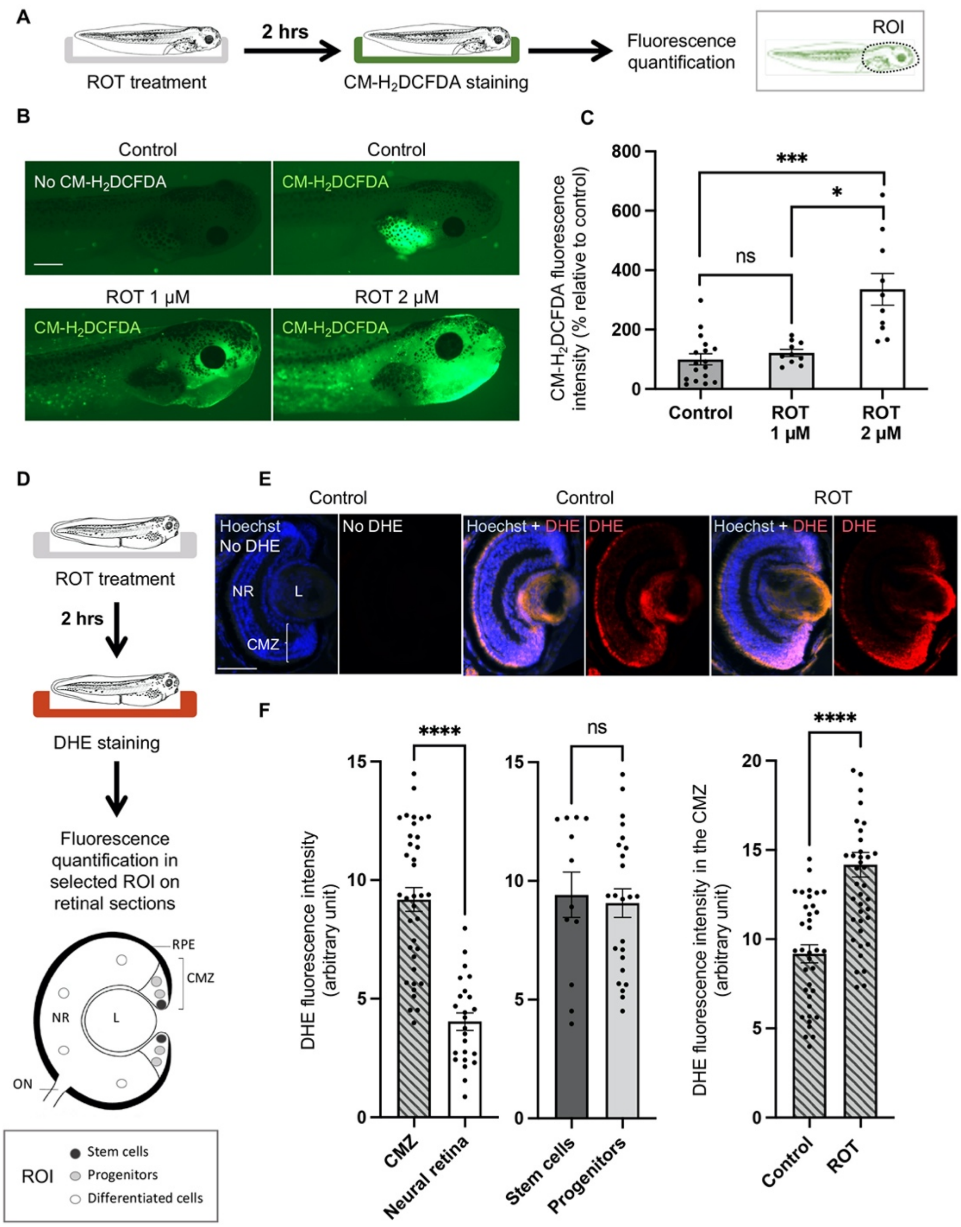


**Supplementary Figure 1 (relative to Figure 1). (A)** Timeline diagram of the experimental procedure used in (B, C). Stage 43/44 control and rotenone (ROT)-treated tadpoles were labelled or not for ROS content with the fluorescent sensor CM-$H_2$DCFDA (CM-$H_2$DCFDA/no CM-$H_2$DCFDA). CM-$H_2$DCFDA intensity was quantified in the tadpole anterior region (delineated in black). **(B, C)** Corresponding images and quantifications. **(D)** Timeline diagram of the experimental procedure used in (E, F) to analyse DHE staining on retinal sections**.** Circles in the retina schematic indicate the 10 regions of interest (ROI) where DHE staining was measured on each section. Dark and light grey circles are located in the stem cell and progenitor regions of the CMZ, respectively. White circles are located within the neural retina, where cells are differentiated. **(E)** Retinal sections from stage 39/40 control and rotenone-treated embryos stained or not with dihydroethidium (DHE/no DHE). Nuclei were counterstained with Hoechst. **(F)** Corresponding quantifications. The left graph includes all measures in ROI located in either the CMZ or neural retina. The central graph distinguishes ROI corresponding either to the stem or progenitor region of the CMZ. The right graph shows pooled measures for all ROI located in the CMZ. Data are represented as mean ± SEM. In (C), 10 to 17 tadpoles were analyzed per condition. In (F), 6 (control) or 7 (ROT) sections were analyzed. Statistics: Mann-Whitney test. CMZ: ciliary marginal zone; L: lens; NR: neural retina; ON: optic nerve; RPE: retinal pigmented epithelium. Scale bars: 500 µm in B and 50 µm in E.

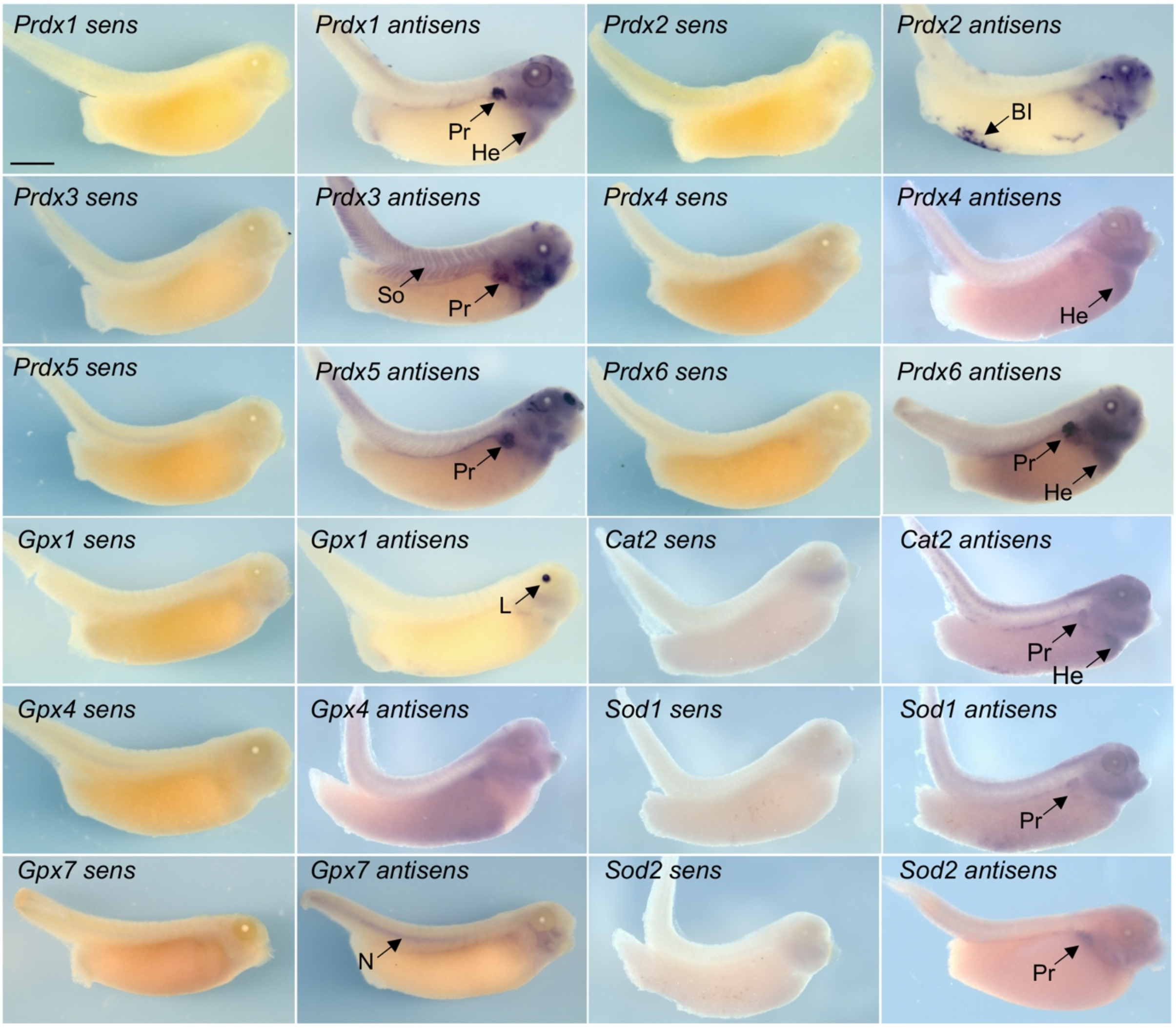


**Supplementary Figure 2 (relative to Figure 2).** Whole mount *in situ* hybridization analysis of antioxidant gene expression on stage 38/39 embryos. Shown are representative images of stainings *in toto* (lateral views) following hybridization with sense or antisens probes. Enlargements of the head regions are shown on Figure 2A. Beyond the head region, some organs displaying high expression level of the considered gene are highlighted (black arrows). BI: blood islands; Cat: catalase; Gpx: glutathione peroxidase; He: heart; L: lens; N: notochord; Pr: pronephros; Prdx: peroxiredoxin; So: somites; Sod: superoxide dismutase. Scale bar: 1 mm.

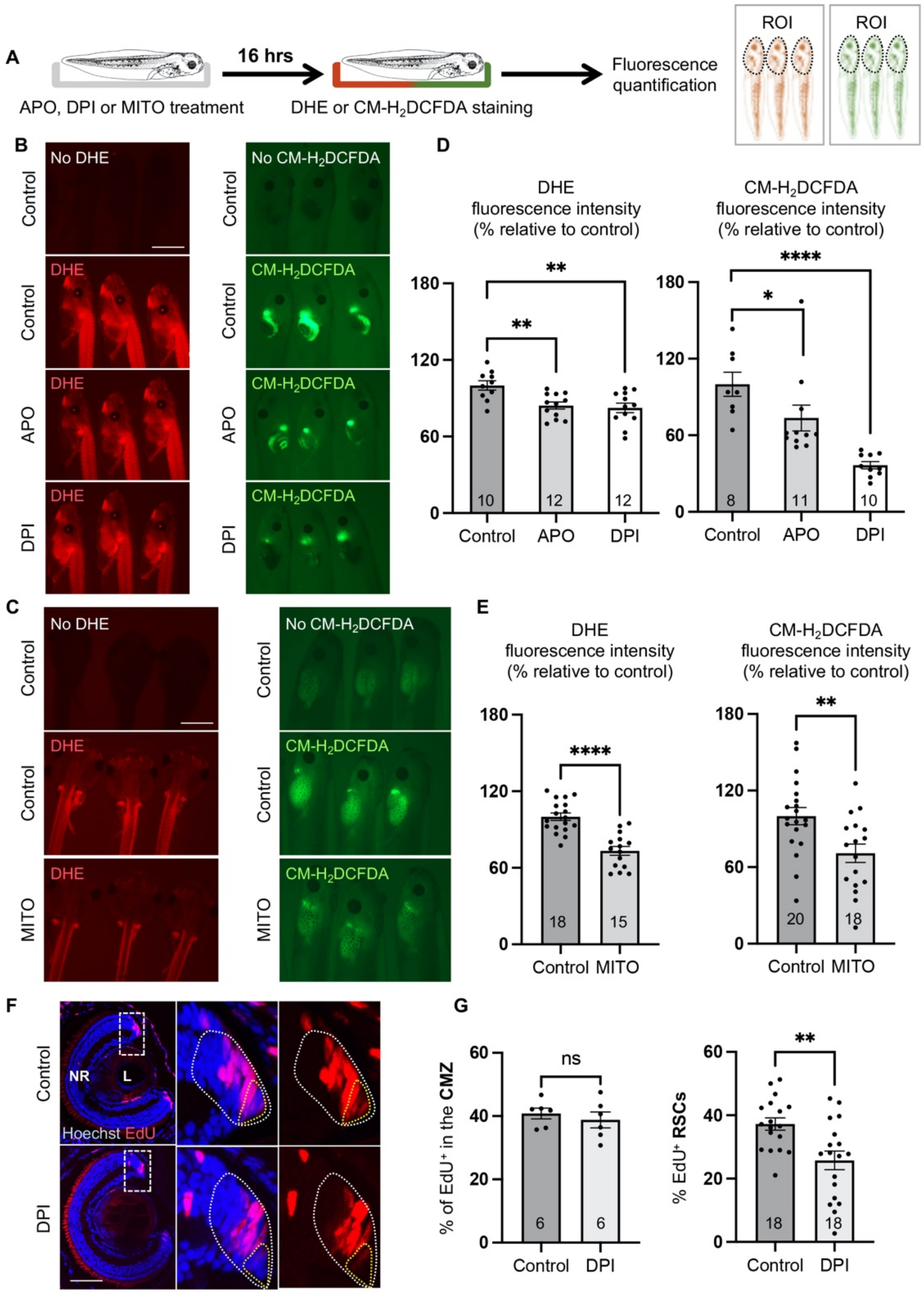


**Supplementary Figure 3 (relative to Figure 3). (A)** Timeline diagram of the experimental procedure used in (B-E). Tadpoles were treated for 16-hours with the NOX inhibitors apocynin or diphenyleneiodonium (APO, DPI; stage 44/45; B), or with the mitochondrial ROS scavenger mitotempo (MITO; stage 42/43; C). They were then labelled for ROS content with the fluorescent sensors dihydroethidium (DHE) or CM-$H_2$DCFDA. Fluorescence intensity was quantified in the anterior region (delineated in black). **(B, C)** Representative images of labelled tadpoles. **(D, E)** Corresponding quantifications. **(F, G)** EdU incorporation assay (4-hour exposure) on retinal sections from stage 43 tadpoles that were treated for 16 hours with diphenyleneiodonium (DPI). Nuclei were counterstained with Hoechst. Right panels are higher magnifications of the dorsal CMZ (delineated in white). RSCs are delineated in yellow. EdU-positive cells were quantified among total CMZ cells or among RSCs. In graphs, data are represented as mean ± SEM. The number of analyzed tadpoles/retinas is indicated in each bar. Statistics: Mann-Whitney test. L: lens; NR: neural retina. Scale bars: 1 mm in B, C and 50 µm in F.

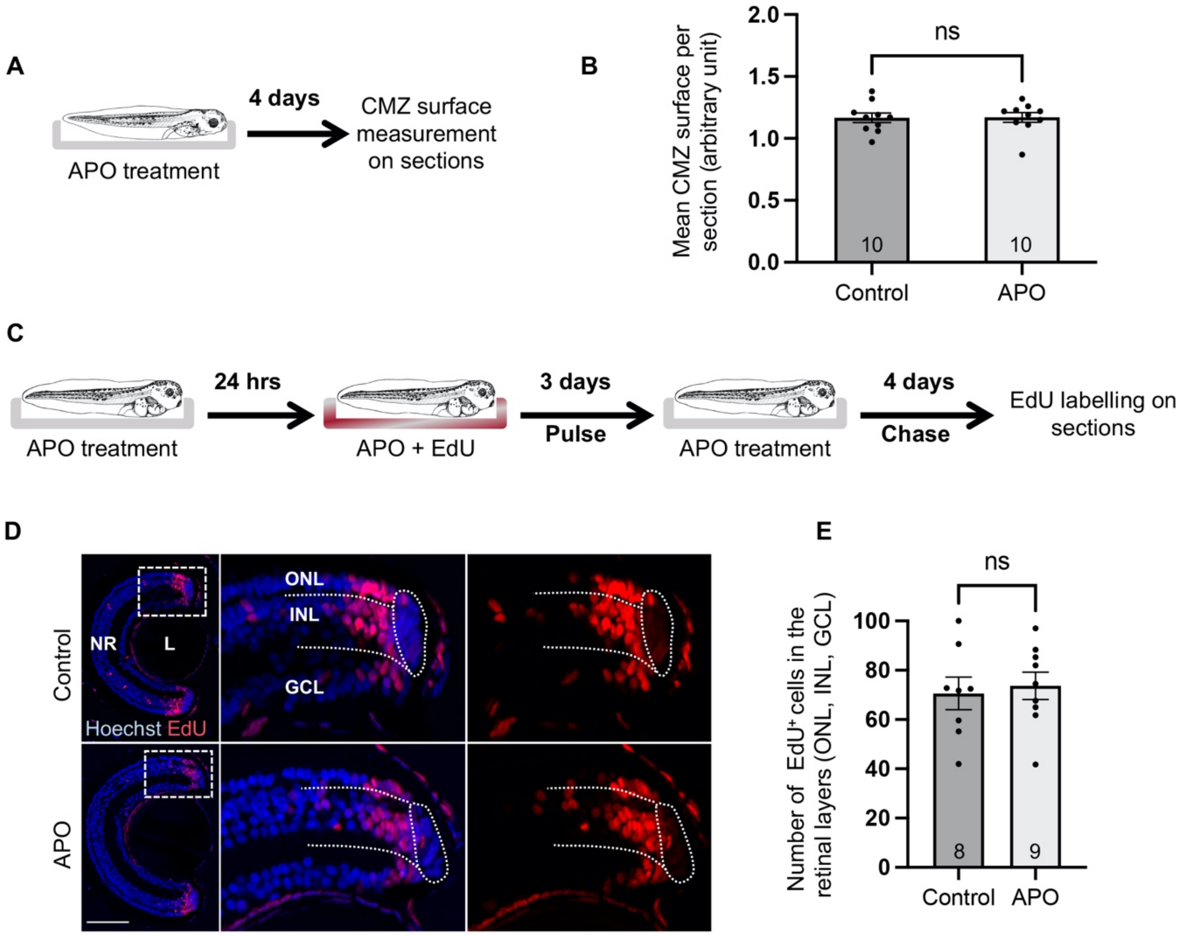


**Supplementary Figure 4 (relative to Figure 3). (A)** Timeline diagram of the experimental procedure used in (B). Stage 42/43 tadpoles were treated for 4 days with apocynin (APO). CMZ surface was then measured on retinal sections. **(B)** Corresponding quantification. **(C)** Timeline diagram of the pulse-chase experiment performed in (D, E). Stage 41/42 tadpoles pre-treated with apocynin (APO) were subjected to EdU exposure for 3 days to label all proliferative cells, and then left for 4 days without EdU. APO was applied to the rearing medium during the whole time-period. **(D, E)** Representative images of retinal sections and corresponding quantifications. EdU-labelled newborn cells were counted in the retinal cell layers. GCL: ganglion cell layer, INL: inner nuclear layer; ONL: outer nuclear layer. In graphs, data are represented as mean ± SEM. The number of analyzed retinas is indicated in each bar. Statistics: Mann-Whitney test.  Scale bar: 50 µm.

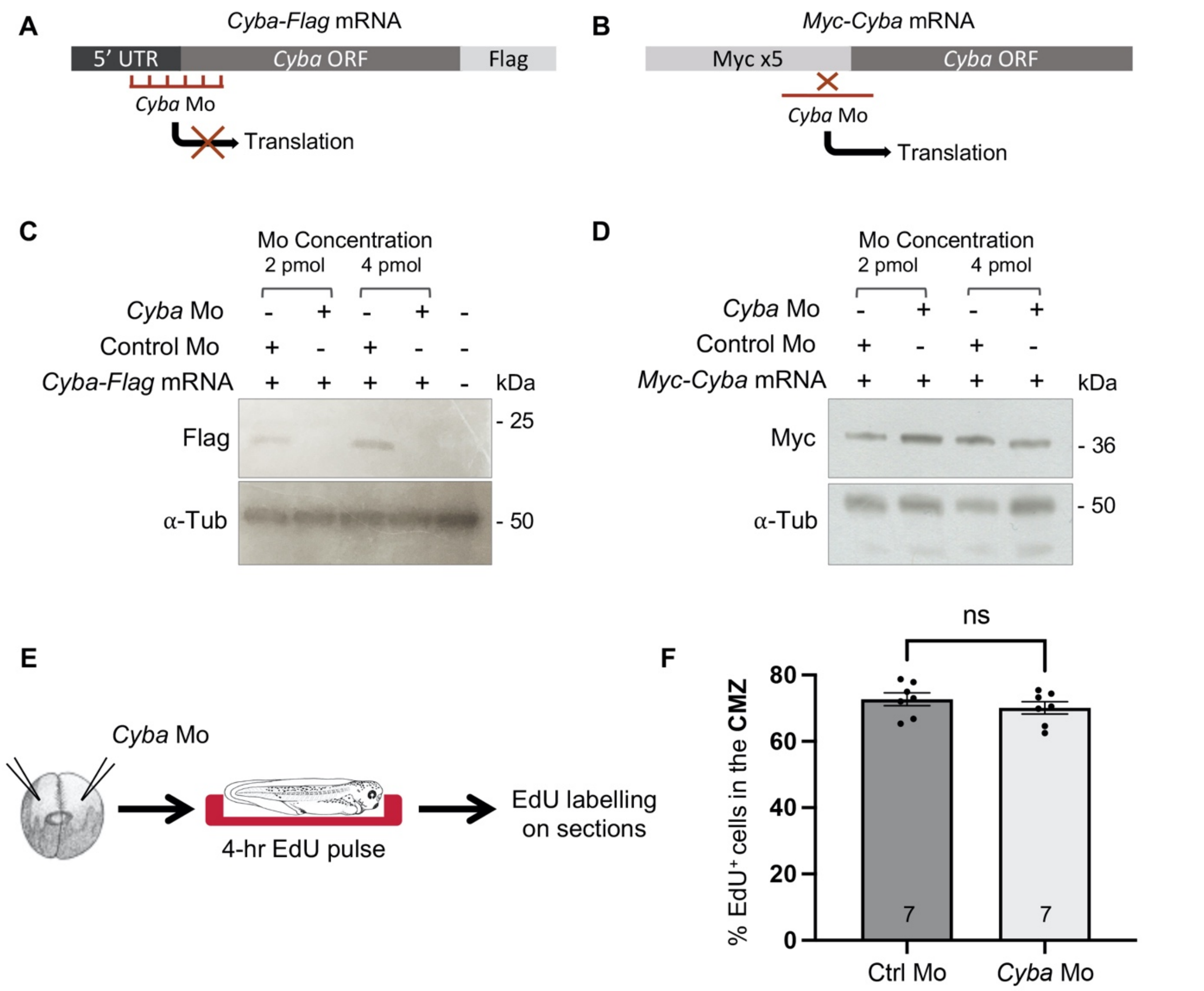


**Supplementary Figure 5 (relative to Figure 3). (A, B)** Schematic representations of *Cyba-Flag* mRNA (reporter construct sensitive to the *Cyba* Morpholino) and *Myc-Cyba* mRNA (rescue construct insensitive to the *Cyba* Morpholino). **(C, D)** Detection of tagged CYBA proteins by Western-blot in whole-embryo extracts, following co-injection at the two-cell stage of control or *Cyba* Morpholinos (Mo), together with either *Cyba-Flag* mRNA (C) or *Myc-Cyba* mRNA (D). Two doses of Morpholinos were assessed (2 or 4 pmol). Proteins were detected with anti-Flag or anti-Myc antibodies. α-Tubulin labelling was used as a loading control. ORF: open reading frame; UTR: untranslated region. **(E)** Timeline diagram of the experimental procedure used in (F). Stage 37/38 embryos were subjected to an EdU incorporation assay (4-hour exposure), following injection at the two-cell stage of control (Ctrl) or *Cyba* Morpholinos (Mo). **(F)** EdU-positive cells were quantified among total CMZ cells in the dorsal region. Data are represented as mean ± SEM. The number of analyzed retinas is indicated in each bar. Statistics: Mann-Whitney test.

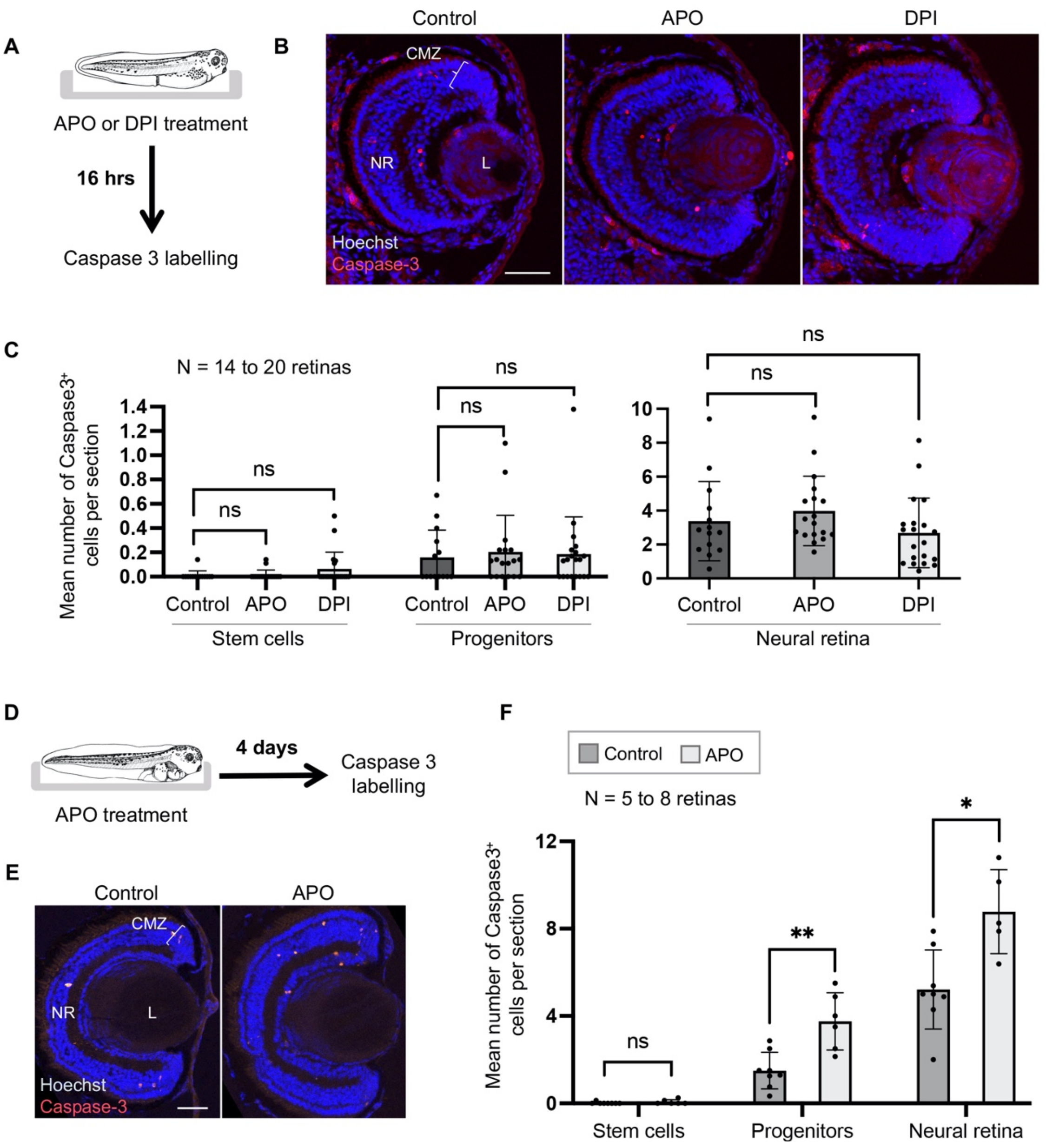


**Supplementary Figure 6 (relative to Figure 4). (A)** Timeline diagram of the experimental procedure used in (B, C). Stage 40 embryos were treated for 16 hours with apocynin (APO) or diphenyleneiodonium (DPI). They were then subjected to immunofluorescence analysis of cleaved Caspase-3 expression. **(B, C)** Representative images of retinal sections and corresponding quantifications. **(D)** Timeline diagram of the experimental procedure used in (E, F). Stage 42 tadpoles were treated for 4 days with apocynin (APO). They were then subjected to immunofluorescence analysis of cleaved Caspase-3 expression. **(E, F)** Representative images of retinal sections and corresponding quantifications. Nuclei were counterstained with Hoechst. Caspase-3-positive nuclei were quantified among RSCs or progenitors of the CMZ and within the neural retina. In graphs, data are represented as mean ± SEM. The number of analyzed retinas is indicated above each graph. Statistics: Mann-Whitney test. CMZ: ciliary marginal zone; L: lens; NR: neural retina. Scale bars: 50 µm.

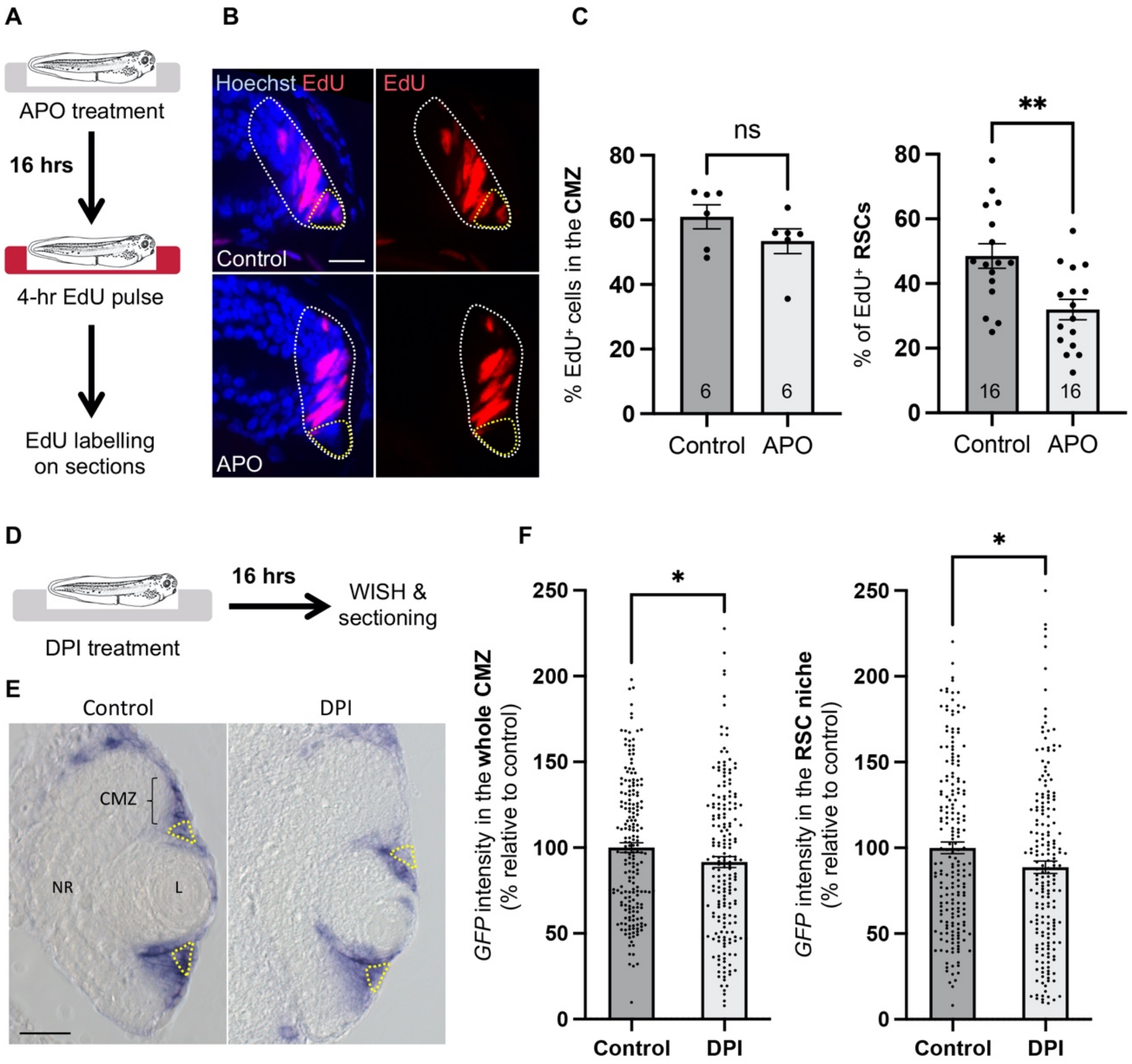


**Supplementary Figure 7 (relative to Figure 7). (A)** Timeline diagram of the experimental procedure used in (B, C). Stage 40 *Xenopus tropicalis* embryos were treated for 16 hours with apocynin (APO). They were then subjected to an EdU incorporation assay (4-hour exposure). **(B)** Representative images of the dorsal CMZ (delineated in white). RSCs are delineated in yellow. Nuclei were counterstained with Hoechst. **(C)** Quantifications of EdU-positive cells among total CMZ cells or among RSCs. Data are represented as mean ± SEM. The number of analyzed retinas is indicated in each bar. **(D)** Timeline diagram of the experimental procedure used in (E, F). Stage 40 *Tg(pbin7Lef-dGFP) Xenopus tropicalis* embryos were treated for 16 hours with diphenyleneiodonium (DPI). They were then subjected to whole mount *in situ* hybridization (WISH) analysis of *GFP* expression. **(E)** Representative retinal sections. The RSC-containing region is delineated in yellow. In the graph, data are represented as mean ± SEM and each point corresponds to a measurement performed either in the dorsal or ventral part of the CMZ. 106 to 112 sections per condition were analyzed (from 42 embryos). Statistics: Mann-Whitney test. CMZ: ciliary marginal zone; L: lens; NR: neural retina. Scale bars: 25 µm in (B), 50 µm in (E).
